# KaryoTap Enables Cost-Effective High-Throughput Single-Cell Aneuploidy Profiling and Reveals How Positive and Negative Selection Shapes Cancer Genomes

**DOI:** 10.1101/2023.09.08.555746

**Authors:** Joseph C. Mays, Fabio Alfieri, Maria Trifas, Manjunatha Kogenaru, Helberth M. Quysbertf, Sally Mei, Nazario Bosco, Xin Zhao, Joy Bianchi, Aleah Goldberg, Gururaj Rao Kidiyoor, Liam J. Holt, David Fenyö, Teresa Davoli

## Abstract

Aneuploidy—the presence of chromosome gains and losses—is highly prevalent in human tumors, yet the contribution of chromosome missegregation rates and selection to shape these patterns remains poorly understood. To address this challenge at scale, we developed KaryoTap, a cost-effective single-cell DNA sequencing method combining custom targeted panels for the Tapestri platform with a Bayesian gaussian mixture model to detect chromosome- and chromosome arm-scale aneuploidy as well as integrated gRNAs and barcodes. KaryoTap achieves an average accuracy of >85% for arm events and >90% for chromosome events at <$1 per cell, allowing scalable analysis of tens of thousands of cells per experiment. Through KaryoTap-based analysis of 11,555 cells, we performed in vitro evolution screens on immortalized human cells from mammary gland, pancreas and melanocytes. By comparing aneuploidy frequencies immediately after reversine-induced missegregation versus after extended proliferation, we quantified positive and negative selection for specific aneuploidies. Most aneuploidies—both gains and losses—are under negative selection, yet some chromosomal gains are under positive selection (such as 8q and 7q). We found that proliferative selective pressures can explain tissue-specific patterns of chromosomal gains observed in human cancers. *Critically, when positively selected events were excluded from analysis, correlations between in vitro and cancer gain frequencies strongly decreased or disappeared, whereas excluding negatively selected events largely preserved these correlations.* These findings demonstrate that proliferative selection shapes the landscape of chromosomal gains in cancer, with a more prominent role for positive selection.

## Introduction

A critical hallmark of cancer initiation and progression is the presence of aneuploidy, or gains and losses of whole chromosomes or chromosome arms, which arise due to mitotic missegregation events(1–3). In addition to aneuploidy, chromosomal instability, characterized by a continuously high rate of these missegregation events among tumor cells, is a driver of tumor progression and metastasis(4,5). Chromosomal instability produces cell populations with heterogeneous aneuploid karyotypes that continuously evolve over time, granting tumor cells an opportunity to adapt to their environment and develop resistance to cancer therapies(6–8). Cancer aneuploidy is characterized by non-random patterns in which specific chromosomes are frequently gained or lost, while others remain largely unchanged. These patterns are often tissue-specific, with certain chromosomal alterations being common in some tumor types but rare or absent in others.

The genomic patterns of aneuploidy in cancer reflect both the frequency at which specific chromosomal alterations occur and the selective pressures that determine their persistence during tumor evolution (58–59, 68). Intrinsic genomic features influence the baseline rate of chromosome missegregation: chromosome size has been proposed to affect the probability of segregation errors, with larger chromosomes potentially experiencing higher rates of missegregation (59). In contrast, selective pressures shape which aneuploidies are retained, as the cumulative effects of oncogenes, tumor suppressor genes, and essential genes on each chromosome determine the fitness consequences of gains and losses in a tissue-specific manner (58, 67). While selection clearly plays an important role in shaping cancer aneuploidy patterns, the relative contributions of positive versus negative selection remain unclear. Recent computational analyses using machine learning approaches have suggested that negative selection may play a greater role than positive selection in shaping these patterns, with tumor suppressor gene density being a better predictor of chromosomal gain patterns than oncogene density (69). However, direct experimental evidence distinguishing between the contributions of missegregation frequency and selective pressures—and between positive and negative selection—in human cells remains limited.

Traditional methods for detecting aneuploidy, such as whole genome sequencing (WGS), rely on bulk averaging of cell populations, effectively masking heterogeneity among individual cells and preventing assessment of the extent or rate of chromosomal instability. Single-cell approaches, particularly single-cell DNA sequencing (scDNA-seq), overcome this limitation(9–13). These data can be used to reassemble the karyotypic evolution of the tumor cells, which can provide insights into how aneuploidy and chromosomal instability drive tumorigenesis and inform treatments for therapeutic resistance(8,11). Methods for modeling aneuploidies of specific chromosomes in cell culture have also emerged as powerful tools for studying the effects of aneuploidy in cancer(14–16). These methods have benefitted from scDNA-seq, as sequencing individual cells enables the evaluation of the specificity and accuracy of the engineered karyotypes(15,16).

Single-cell DNA sequencing (scDNA-seq) methods have emerged as powerful tools for detecting aneuploidy in individual cells, though each approach involves tradeoffs between throughput, sensitivity, and cost (17–24). For scDNA-seq methods based on untargeted whole-genome sequencing (WGS), there is an inherent bias in the sensitivity of detecting whole-chromosome or arm-level gains and losses. Because sequencing reads are distributed proportionally across the genome, smaller chromosomes receive fewer reads than larger ones and are thus detected with lower sensitivity. However, when quantifying aneuploidy or chromosomal instability - defined as the rate of chromosome missegregation – each chromosome is counted as a single unit regardless of its size. Therefore, to achieve uniform sensitivity across all chromosomes, WGS-based scDNA-seq requires high sequencing depth, which increases cost substantially.

To address the need for a high-throughput, high-sensitivity, cost-effective method for detecting chromosome-scale aneuploidy across the human genome, we turned to the Tapestri platform from Mission Bio, a droplet-based targeted scDNA-seq solution that allows for the sequencing of hundreds of genomic loci across thousands of cells in one experiment(25). The platform is commonly used to detect tumor hotspot mutations in cancer driver genes, yet it has not been utilized to identify aneuploidy across the genome. While targeted sequencing is typically used to identify single nucleotide mutations(26), we reasoned that we could use the relative sequencing depth of targeted loci to detect chromosome- and chromosome arm-scale aneuploidy in individual cells. Here, we describe KaryoTap, a cost-efficient method combining custom targeted Tapestri panels of PCR probes covering all human chromosomes with a Gaussian mixture model framework for DNA copy number detection, thus enabling the accurate detection of aneuploidy in thousands of heterogeneous cells. Additionally, we included probes that detect lentiviral-integrated CRISPR guide RNAs to enable functional studies and DNA barcodes to enable sample multiplexing. This method also enables detection of copy number neutral loss of heterozygosity (CNN-LOH) events due to its high sequencing depth. To enhance usability, we also developed a companion software package for R, *karyotapR*, which enables the streamlined copy number analysis, visualization, and exploration of the data produced by our custom panels. By overcoming the limitations of current methods, this system will be a valuable tool for investigating the evolution and consequences of aneuploidy and chromosomal instability in human tumors, in addition to other healthy and diseased tissues.

Using KaryoTap, we conducted *in vitro* single-cell proliferation-based karyotype screens using different types of immortalized untransformed human cells. Following treatment with an MPS1 inhibitor to induce chromosomal instability and random aneuploidies, we allowed the cells to proliferate in culture over several weeks. We observed that the patterns of chromosomal gains that emerged during *in vitro* proliferation closely mirrored the aneuploidy patterns observed in human cancers. In contrast, the patterns of chromosomal losses seen in tumors were not recapitulated *in vitro,* suggesting that the *in vivo* tissue microenvironment may play a critical role in shaping these specific aneuploidy events *in vivo*.

These experiments demonstrate that Karyo-Tap is a powerful approach for uncovering the selective forces that shape karyotype evolution in cancer and other biological contexts.

## Results

### Design of a Custom Targeted Panel for Detecting Chromosome-Scale Aneuploidy

To enable accurate, cost-effective detection of aneuploidy at scale in thousands of single cells, we designed custom targeted panels for the Tapestri system. Our Version 1 panel (v1; **Fig 1**; CO261) is comprised of 330 PCR probes that target and amplify specific loci across all 22 autosomes and the X chromosome (**Fig 1B**; **Table S1**; **Additional File 1**). The number of probes targeting each chromosome was proportional to chromosome size (e.g., 24 probes for chr1, 5 for chr22) to achieve a roughly uniform density of ∼1-2 probes per gigabase across all chromosomes. Loci were selected to cover regions carrying single nucleotide polymorphisms (SNPs) with major allele frequencies of 0.5-0.6 such that cells from different lines or individuals sequenced in the same experiment could be identified by their distinct genotypes. This is a key feature of our methodology compared to other scDNA-seq methods, as it allows detection of loss of heterozygosity (LOH) including copy number-neutral LOH.

**Figure 1.**
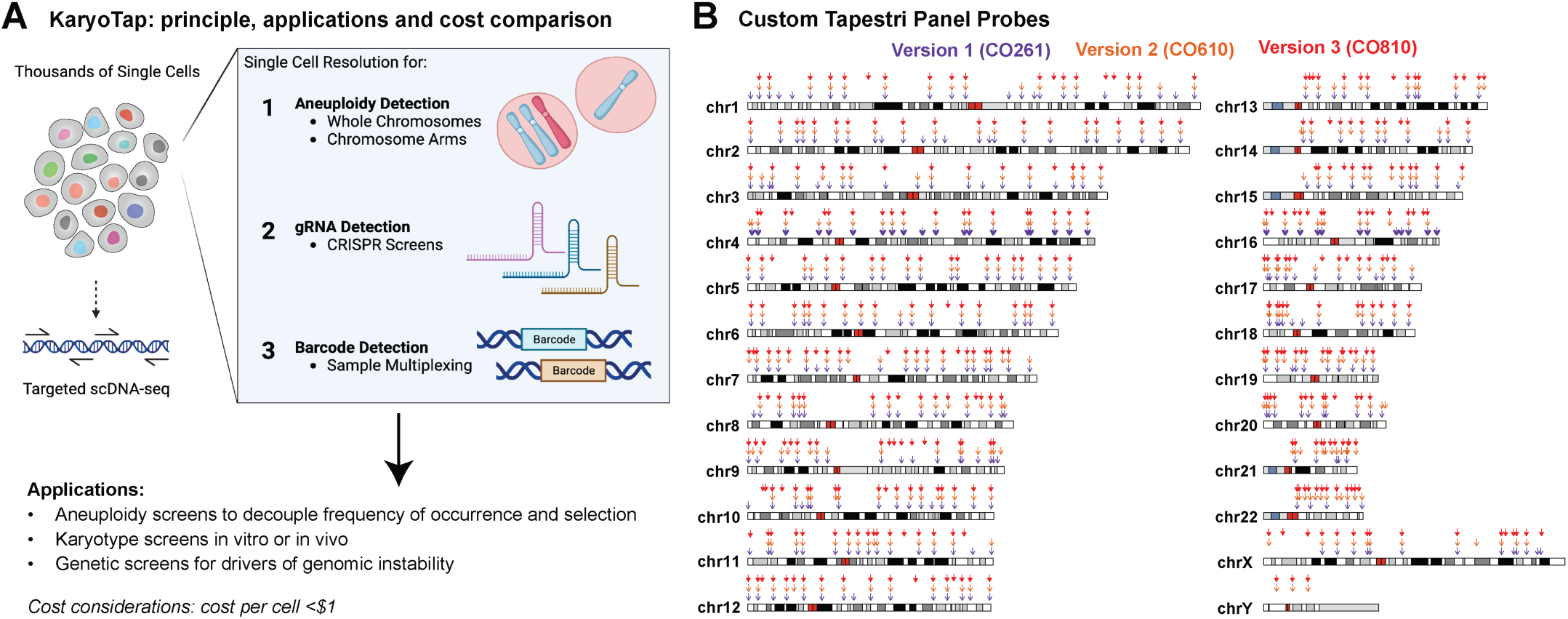
(A) Summary of KaryoTap as a novel single cell DNA sequencing method for high-throughput detection of aneuploidy in single cells in a cost efficient manner. **(B)** Map of PCR probe locations (arrows) for custom Tapestri panels for KaryoTap Version 1 (CO261), Version 2 (CO610) and Version 3 (CO810) on human genome hg19. Red blocks indicate centromeres, grayscale blocks indicate the G-band intensity of cytobands, and blue blocks indicate acrocentric chromosome arms.

### Population-Level Validation of KaryoTap Copy Number Estimates

To validate that KaryoTap accurately detects chromosomal copy numbers, we pooled five cell lines with distinct karyotypes and analyzed 2,986 cells in a single experiment. The pool consisted of near-diploid RPE1 (+10q, XX)(6,27) as a reference population and four aneuploid colon cancer cell lines: LS513 (+1q, +5, +7, +9, +13, +13, XY), SW48 (+7, +14, XX), LoVo (+5p, +7, +12, XY), and CL11 (-6, +7, +7, -17, -18, -22, XY). Known karyotypes were confirmed by bulk WGS (**Fig S1A**) and G-banded karyotyping for RPE1 (**Fig S1B**).

Dimensionality reduction of variant allele frequencies (VAFs) by PCA and UMAP revealed 5 distinct clusters corresponding to each cell line (**Fig S1C**), demonstrating that multiplexed samples can be resolved by genotype alone. We identified the RPE1 cluster by comparing mean VAFs to published deep WGS data(28) (**Fig S1D-E**) and used these cells as the diploid reference for copy number estimation. Cell-probe copy number scores were calculated as the ratio of normalized read counts to the median of the RPE1 reference population, then smoothed across probes targeting each chromosome to generate cell-chromosome scores (see Methods).

At the population level, copy number profiles matched the expected karyotypes from bulk WGS for all five cell lines (**Fig S2A**). For example, averaged heatmaps correctly showed 3 copies of chromosome 7 in LS513, SW48, and LoVo, and 4 copies in CL11; monosomy of chromosome 6 in CL11; and single X chromosomes in male cell lines. Importantly, unbiased clustering distinguished all five populations by copy number alone (**Fig S2B**), mimicking the subclonal structure of a heterogeneous tumor(31). Further clustering within LoVo cells revealed two subpopulations distinguished by exclusive gains in chr5p or chr15q (**Fig S2C**), demonstrating KaryoTap’s ability to detect clonal heterogeneity. While this confirms KaryoTap’s accuracy at the bulk level, accurate copy number calling in individual cells requires accounting for cell-to-cell variability, which we address below.

### Bayesian Gaussian Mixture Modeling Enables Accurate Integer Copy Number Calling in Single-Cell DNA Sequencing Data

To account for intracellular variation in the distributions of copy number scores, we classified the scores as integer copy number calls using a 5-component Gaussian mixture model (GMM)(30), where each component represents a possible copy number value of 1-5 (**Fig S3A**). To train the GMM model, we used the known copy number state of each chromosome in RPE1 cells and simulated the expected distribution of copy number scores for each integer copy number (1–5) separately for each chromosome. These 5 simulated distributions represent the score profiles that would be observed if a chromosome had 1, 2, 3, 4, or 5 copies, and they serve as the component distributions for the Gaussian mixture model. We then applied Bayes theorem to calculate the posterior probability of each cell-chromosome score belonging to each of the 5 GMM component distributions and assigned the cell-chromosome an integer copy number corresponding to the component with the highest posterior probability (**Fig. S3A**; see Methods).

To evaluate copy number calling performance, we determined the accuracy of the classifier model, calculated as the proportion of calls that were correct (i.e., true positives), using the known copy numbers for RPE1 as ground truth on data generated using **KaryoTap Version 1** Panel. We focused on RPE1 as its karyotype is stable and homogeneous across the population(6); karyotyping confirmed that the line is triploid for chr10q and diploid for all other chromosomes in all metaphases analyzed, with the exception of 3 copies of chr12 in 3% of metaphases (**Fig S1B**). Accuracy, or correctly identifying 2 copies of a chromosome in RPE1, varied between chromosomes and ranged from 95% for chr2 to 49% for chr22 with a mean of 79% (**Fig S3B**). Five-fold cross-validation confirmed robustness of these estimates (mean absolute deviation: 0.61 to 4.75 percentage points across chromosomes). Eighteen out of 22 chromosomes had sensitivities of at least 75%; chr10 was excluded from whole-chromosome analysis because the p and q arms have different copy numbers. Accuracy for each chromosome correlated strongly with chromosome length (Pearson r = 0.73), and chromosome length itself correlated strongly with the number of probes targeting the chromosome (Pearson r = 0.93) (**Fig S3C**), suggesting that classifier accuracy is related to the number of probes targeting a chromosome. As expected, linear regression of accuracy on the number of probes per chromosome indicated that the number of probes is predictive of copy number call accuracy (R² = 0.76; p = 1.12e-08) (**Fig S3D-E**). We next performed a similar analysis for chromosome arms (**Fig S4A**). We called integer copy numbers using a GMM generated for each chromosome arm and evaluated classifier accuracy for correctly calling copy numbers in RPE1. The arm-level accuracy ranged from 91% (chr2q) to 50% (chr19p, chr22q) with a mean of 69% (**Fig S4B**). Twenty-two out of 41 arms had accuracy values of at least 70%. A positive relationship between the number of probes and accuracy was again confirmed by linear regression (R² = 0.59; p = 5.1e-09) (**Fig S4C**). Overall, our system accurately calls copy numbers for the majority of chromosomes and chromosome arms, with lower accuracy for smaller chromosomes and arms and that classifier accuracy could be improved by adding additional probes (see also **Supplemental text**).

### Impact of Probe Number and Sequencing Depth on Accuracy

We performed extensive simulation-based analyses to characterize the parameters influencing copy number classification accuracy; full details are provided in the **Supplemental Text** and summarized here. Briefly, probe downsampling experiments on chromosomes 2 and 6 confirmed that probe number, not chromosome size, drives classification accuracy, with median accuracy decreasing substantially as probes were reduced from full coverage to 4 probes per chromosome (**Fig S2D**). Variability in accuracy also increased markedly with fewer probes (**Fig S3A**). Panel simulations using 120 probes from chromosomes 1–6 demonstrated that mean theoretical sensitivity increases with probe number across all copy number levels (**Fig S2E**), achieving ≥90% sensitivity at 4 probes for losses, 20 for diploid states, and 50 for 3-copy gains. A simpler 3-component GMM (loss/neutral/gain) achieved ≥90% sensitivity at 26 probes for gains, with 99% sensitivity requiring 76 probes (**Fig S3B**). In contrast, sequencing depth had modest effects: downsampling from 85 to 17 reads per cell per probe decreased sensitivity by only 4.4 percentage points (**Fig S2F**), indicating that probe number remains the dominant factor while depth beyond ∼35 reads provides minimal additional benefit.

### KaryoTap Version 3 Panel Increases Accuracy of Aneuploidy Detection to ∼90% Keeping the Cost per Cell at a Minimum

As the accuracy of our custom panel increases with the number of probes targeting a chromosome, we optimized our design by adding probes to chromosomes with lower coverage. To balance cost with meaningful sensitivity gains, we removed the 61 least efficient probes (by total read counts) and added 82 probes such that each chromosome was targeted by at least 12 probes (**Table S1**). We also included 4 probes targeting chrY, enabling detection of all 24 chromosomes (**Fig S6A**). The new panel, Version 2 (v2; CO610) comprises 352 total probes, 270 of which are shared with v1 (**Fig 1B**; Additional File 2). To further improve arm-level accuracy, we modified v2 to generate **Version 3** (v3; CO810; **Fig 1B**; Additional File 3). V3 includes 399 total probes, 309 shared with v2, prioritizing chromosome arms with lower coverage. The barcode and gRNA-detecting probes were retained in both designs.

To evaluate v2 and v3 performance, we performed a Tapestri experiment using RPE1 cells (**Fig S6B**) and determined empirical sensitivity as the proportion of cells with correct copy number calls. Five-fold cross-validation confirmed robustness (MAD: 0.49 to 4.74 pp for v2; 0.60 to 3.31 pp for v3). The poorest performing chromosome in v1, chr22, had sensitivity of 62% with v2 (**Fig S6C**) and 78% with v3 (**Fig 2A**), compared to 46% for v1 (**Fig S3B**). Mean sensitivity across all chromosomes was 85% for v2 (**Fig S6C**) and 91% for v3 (**Fig 2A-B**), increased from 79% for v1. Accuracy increased by 2.2 pp on average for each additional probe (**Fig 2C**). Similarly, arm-level average sensitivity increased from 69% (v1) to 76% (v2; **Fig S6D**) to 83% (v3; **Fig 2D-E**), with 2.9 pp improvement per additional probe. Theoretical sensitivity also improved across panels for both 5-component and 3-component GMMs (**Fig S7A**). The decreased noisiness in heatmaps of GMM copy number calls across v1, v2, and v3 for RPE1 (**Fig 2F**), LoVo (**Fig 2G**), and LS513 (**Fig 2H**) further supports improved accuracy with v3. Altogether, panel v3 delivers accurate copy number calls in thousands of single cells with arm-level resolution.

**Figure 2.**
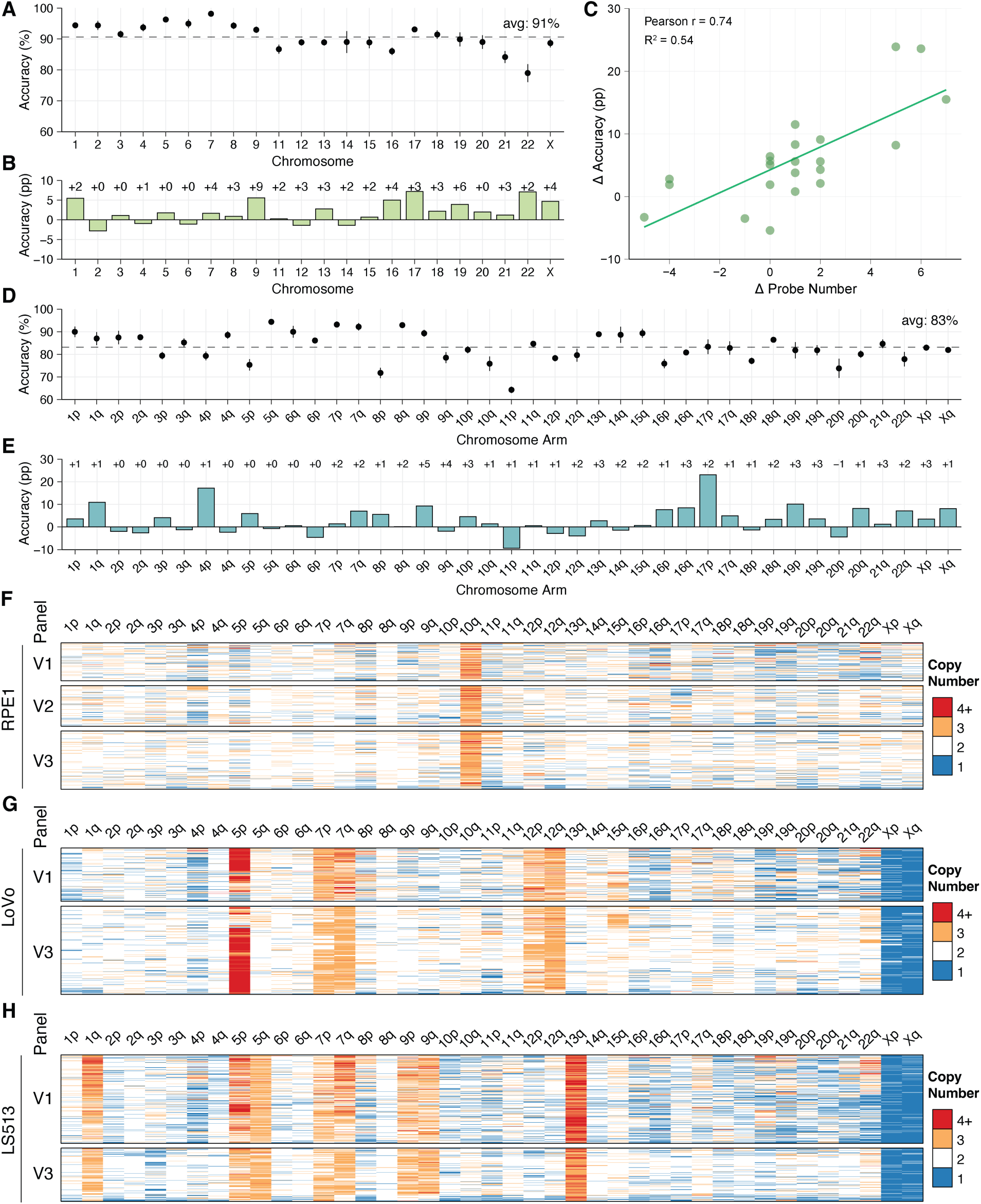
Copy number calling and lentiviral barcoding for custom KaryoTap panel Version 2. **(A)** 5-fold cross validation of copy number call sensitivity for n=631 RPE1 cells by chromosome using panel Version 3. Chr10 is omitted. Dot indicates mean accuracy and lines indicate ± mean absolute deviation. Horizontal dotted line indicates average (avg) accuracy across chromosomes. **(B)** Change in copy number call accuracy by chromosome for RPE1 cells between Version 3 and Version 2 panels. Δ Probe Number is the difference in number of probes targeting a given chromosome between Version 3 and Version 2. pp: percentage points. **(C)** Linear regression of the change in copy number call accuracy for RPE1 cells between Version 3 and Version 2 panels on the change in probe number targeting each chromosome. **(D)** 5-fold cross validation of copy number call accuracy for RPE1 cells by chromosome arms using custom Tapestri panel Version 3. Dot indicates mean accuracy and lines indicate ± mean absolute deviation. Horizontal dotted line indicates average accuracy across chromosome arms. **(E)** Change in copy number call sensitivity by chromosome arm for RPE1 cells between Version 3 and Version 2 panels. **(F-H)** Heatmaps of GMM copy number calls for chromosome arms using panels Version 1, 2, and 3 for cell lines RPE1, LoVo, and LS513.

The main purpose of KaryoTap is to detect chromosomal instability and aneuploidy at the chromosome level. In scDNA-seq methods based on untargeted WGS (e.g., Arc-well, 60), sensitivity of aneuploidy detection depends on total reads per chromosome and thus is inherently lower for smaller chromosomes. For evaluating aneuploidy and chromosomal instability, each chromosome counts as a single entity regardless of size. Thus, WGS-based scDNA-seq methods require high read depth—and higher cost—to achieve sufficient sensitivity across all chromosomes. KaryoTap experiments cost approximately $0.5-1 per cell—less than 10% of alternative methods (see also Supplemental Text). By achieving comparable detection sensitivity at a fraction of the cost compared to WGS methods, KaryoTap provides a scalable and cost-effective solution for high-throughput aneuploidy detection that overcomes the size-dependent biases inherent to WGS-based approaches.

### Detection of Lentiviral Barcodes and gRNAs Allows for Genetic Screenings: Example of KaryoCreate-based Screen for Centromeric sgRNAs

To extend the capabilities of our system for functional genomics, we added two probes to the Version 2 panel that target and amplify either a DNA barcode sequence or CRISPR guide RNA (gRNA) sequence integrated into a cell’s genome by lentiviral transduction. DNA barcoding can be used when several samples from the same cell line or individual are sequenced together and cannot be distinguished by genotype. Similarly, CRISPR gRNAs can be used both for functional studies and as barcodes themselves, indicating the gRNA treatment received by each individual cell.

As proof-of-concept, we transduced RPE1 cells and human colorectal epithelial cells (hCECs) each with distinct gRNA constructs (gRNA1 and gRNA2, respectively), and human Pancreatic Nestin-Expressing cells (hPNEs) with a mix of two DNA barcode constructs that drive BFP expression. We used distinct cell lines for each construct so the three populations could be distinguished by genotype without assuming successful barcoding. Probe AMP350 targets the region surrounding and including the gRNA sequence in the lentiviral vector; Probe AMP351 targets the barcode sequence region (**Fig 3A**).

**Figure 3.**
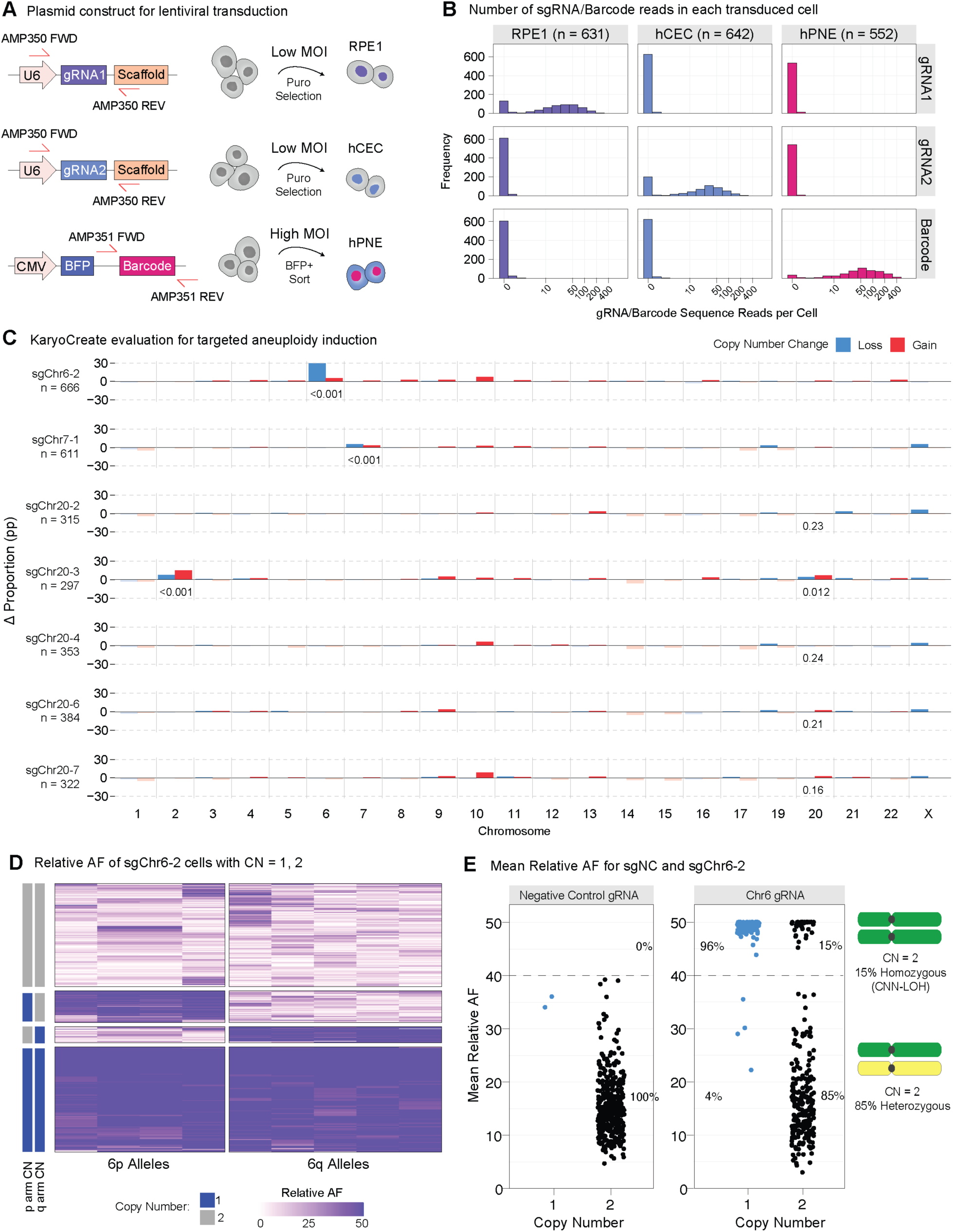
Evaluation of KaryoCreate and Loss of Heterozygosity. **(A)** Plasmid constructs for lentiviral transduction of RPE1, hCEC D29, and hPNE cell lines. Probe pair AMP350 amplifies a 253 bp region including a CRISPR gRNA sequence and part of the U6 promoter and F+E scaffold. Probe pair AMP351 amplifies a 237 bp region including a barcode sequence. **(B)** Number of reads in each cell that match the expected sequence of gRNA1, gRNA2, or barcodes in 3 transduced cell lines. X-axis is log-scaled. Number of cells in each cell line is indicated. **(C)** Evaluation of KaryoCreate technology by KaryoTap panel Version 2. Bars represent the percentage point (pp) change (Δ) in the proportion of chromosomal losses (1 copy) and gains (3+ copies), compared to sgNC negative control (n = 398 cells). Chr18 was omitted due to additional copies of the chromosome in the cell line. p-values from Fisher’s Exact test comparing the proportion of cells with copy number = 2 to copy number = {1, 3, or 4} (i.e., diploid vs. aneuploid) in each chromosome in each sample to the corresponding chromosome in the sgNC negative control sample are shown where p < 0.1. Additional p-values > 0.1 are reported where relevant. All p-values are reported in Table S2. Bars with negative values and p > 0.1 have reduced opacity for clarity. **(D)** Heatmap of relative VAFs for sgChr6-2 treated cells with 1 or 2 copies of chr6 as called by GMM, for 9 originally heterozygous variants. Relative allele frequencies calculated as the absolute difference between raw allele frequency and 50%. 0 corresponds to balanced heterozygous alleles and 50 corresponds to fully homozygous alleles. Heatmap rows split by k-means clustering where k=4 and sorted by hierarchical clustering. CN: copy number. **(E)** Mean relative VAFs for sgNC and sgChr6-2 treated cells with 1 or 2 copies of chr6 as called by GMM. 0%-40% AF indicates heterozygous haplotype, 40%-50% AF indicates homozygous haplotype.

gRNAs were transduced at a multiplicity of infection (MOI) of 1-1.5 with puromycin selection; BFP barcodes were transduced at higher MOI with FACS enrichment for BFP expression. The three populations were pooled and analyzed in a single Tapestri experiment using panel Version 2, with populations distinguished by PCA, UMAP, and clustering of VAFs (**Fig S6B**). RPE1 cells had an average of 34 gRNA1 reads per cell (0 in other lines), hCECs had 30 gRNA2 reads per cell, and hPNEs had 81 barcode reads per cell (**Fig 3B**). Detection rates were 79% for RPE1 gRNA1, 69% for hCEC gRNA2, and 94% for hPNE barcodes (**Fig 3B**), with virtually all probe reads matching expected sequences (**Fig S7B**), indicating high specificity with no contamination. Detection rates can be further improved through higher MOI transduction or increased sequencing depth.

To demonstrate the combined copy number detection and multiplexing capabilities of our system, we tested it on samples treated with KaryoCreate (Karyotype CRISPR Engineered Aneuploidy Technology), our method for inducing chromosome-specific aneuploidy in cultured cells(14). KaryoCreate uses CRISPR gRNAs to target a mutant KNL1-dCas9 fusion protein to the centromere of a specific chromosome, causing missegregation in ∼20% of cells. KaryoTap represents an ideal method to evaluate the efficiency and chromosome-specificity of KaryoCreate. We performed a Tapestri experiment on hCECs treated with gRNAs previously tested using KaryoCreate: sgNC (untargeted negative control), sgChr6-2 (targets chr6), and sgChr7-1 (targets chr7). The gRNA sequences amplified by AMP350 were used to identify the gRNA each cell received.

sgChr6-2 and sgChr7-1 induced gains and losses specifically in intended chromosomes, but not others, compared to sgNC (**Fig 3C**, **Table S2**; p < 0.001, Fisher’s exact test). sgChr6-2 induced 26.6% losses of chr6 compared to 0.5% with sgNC, and 8.3% gains compared to 2.0% (**Table S6**). sgChr7-1 induced 6.1% losses of chr7 compared to 0.5% with sgNC, and 5.1% gains compared to 4.3%.

In the same experiment, we screened 5 sgRNAs targeting chromosome 20 (sgChr20-2, 20-3, 20-4, 20-6, and 20-7) that were previously described but not validated by imaging (14). Chromosome 20 is one of the most frequently gained chromosomes in human cancer(2), but its small size and low number of centromeric gRNA binding sites (∼700) had prevented validation by imaging or scRNA-seq. sgChr20-2, 20-4, 20-6, and 20-7 did not induce changes in chr20 (p = 0.557-0.926). However, sgChr20-3 induced 11.1% gains in chr20 compared to 6.0% with sgNC, and 5.7% losses compared to 4.0% (p = 0.012). Notably, sgChr20-3 also induced 9.4% losses and 18.5% gains in chr2 (p < 0.001). The sgChr20-3 sequence (GGCAGCTTTGAGGATTTCGT) matches 18 out of 20 base pairs for loci on the chr2 centromere (GATAGCTTTGAGGATTTCGT)(14), suggesting an explanation for this off-target effect.

These data indicate that KaryoTap successfully enables simultaneous detection of aneuploidy and gRNA/barcodes in the same cells and thus can be used to perform CRISPR-based (i.e., gRNA-based) or ORF-based (barcode-based) functional screens.

### Karyotap Enables Detection of Copy Neutral Loss of Heterozygosity (CNN-LOH)

Gains and losses of diploid chromosomes result in a shift in VAF for their heterozygous SNPs from 50% in the direction of 100% or 0% depending on which parental chromosome copy (i.e., haplotype) experienced a copy number change. In addition, a shift in VAF can also be observed in the absence of copy number changes in copy number neutral loss of heterozygosity (CNN-LOH), which has been observed in cancer as well as normal tissues. Because each probe is sequenced at a high depth, KaryoTap is able to detect this shift, allowing us to determine which of the two parental chromosomes/haplotypes was gained or lost. This is especially important for detecting loss of heterozygosity (LOH), a common event in cancer whereby a heterozygous-to-homozygous shift by chromosomal loss can inactivate tumor suppressor genes(31,32, 71). To determine if KaryoTap could detect allele frequency shifts following chromosomal gains and losses, we examined the cells from the KaryoCreate experiment (**Fig 3A**) that had lost a copy of chromosome 6 after treatment with the sgChr6-2 gRNA. We identified 9 heterozygous variants on chr6 by identifying variants with a mean allele frequency between 20-80% in the sgNC control population. We then calculated a relative (i.e., haplotype-agnostic) allele frequency for sgChr6-2 treated cells with 1 or 2 copies of chr6 by calculating the absolute difference between raw allele frequency and 50% such that 0 corresponded to heterozygous alleles and 50 corresponded to fully homozygous alleles (**Fig 3D**). 4 distinct clusters of cells emerged using K-means clustering on relative AFs. The cluster comprising cells with a copy number call of 2 for chr6 shows that the 9 variants are heterozygous in diploid cells as expected. The cluster with one copy of chr6 shows a shift across the variants from heterozygous to homozygous (i.e. a loss of heterozygosity), supporting the loss of one copy of each allele in these cells. There are also two smaller clusters representing the loss of either chromosome arm but not the other, supported by both the loss of heterozygosity in the variants on the affected arm and the copy number call of 1 for that arm (**Fig 3D**). This indicates that KaryoTap can be used to detect loss of heterozygosity in single cells at the population level.

It is possible that the loss of a chromosome can be followed by a duplication of the remaining chromosome, such that the copy number of the chromosome remains the same, but one allele is lost, i.e., a CNN-LOH. To determine if KaryoCreate can cause CNN-LOH in the targeted chromosome, we took the relative allele frequencies for sgChr6-2 treated cells with 1 or 2 copies of chr6 calculated above and averaged them such that each cell had one mean relative AF value for chr6. We also repeated this calculation for the cells treated with the sgNC control gRNA. If we consider relative AF between 40% and 50% to indicate homozygosity of chr6 alleles and 0% to 40% to indicate heterozygosity, all sgNC-treated cells with a chr6 copy number of 2 were heterozygous (**Fig 3E**). Cells that lost a copy of Chr6 following treatment with sgChr6-2 were detected to be homozygous for their remaining allele. 85% of cells treated with sgChr6-2 that had 2 copies of chr6 detected were heterozygous, while 15% were detected as homozygous, indicating a loss of heterozygosity with no change in the net copy number in this population (2 copies of chr6). This indicates that KaryoCreate can induce CNN-LOH, and KaryoTap can detect CNN-LOH events.

### KaryoTap enables single cell-based screens to differentiate between occurrence rate and selection among chromosomal gains and losses

We next utilized KaryoTap to perform a karyotype evolution screening with the goal of assessing whether the gain or loss of any chromosome or chromosome arm would confer a fitness advantage or disadvantage during cellular proliferation *in vitro*. Given its resolution and scalability, KaryoTap is well-suited for monitoring how aneuploidy patterns occur and are selected under proliferative pressure. To model aneuploidy induction and selection , we exposed nontransformed, near-diploid cell lines to Reversine (an MPS1 inhibitor) to induce chromosomal missegregation and thus gains and losses of all chromosomes (74) (**Fig 4A**). By profiling cells immediately after Reversine exposure and again after several weeks in culture, we monitored how initial random aneuploidy events are progressively filtered by positive and negative selective pressures.

**Figure 4.**
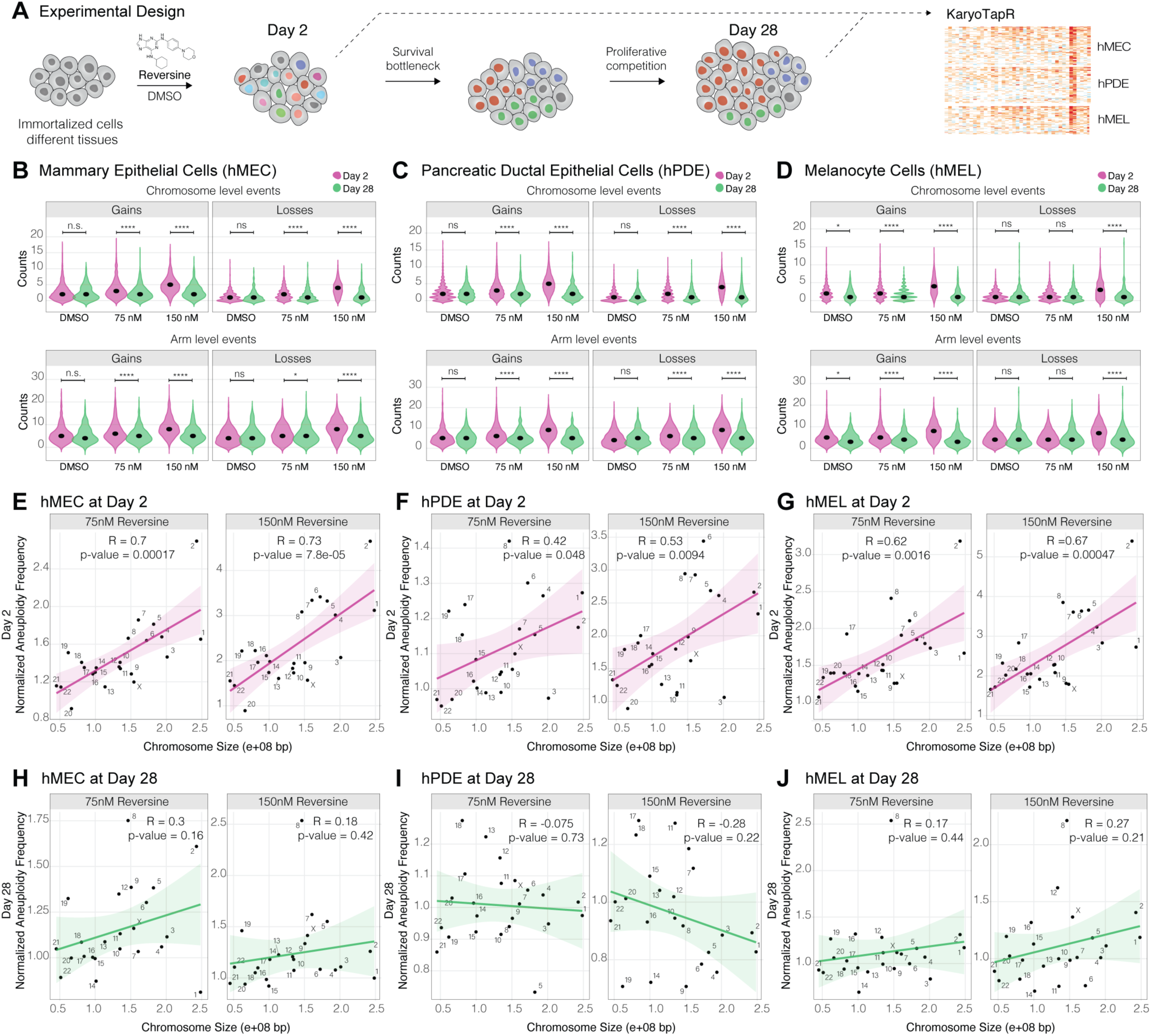
Reversine-Induced chromosomal instability and in vitro selection dynamics. **(A)** Experimental design of reversine-induced chromosomal instability. Healthy cell lines (mammary epithelial cells, hMEC; pancreatic epithelial cells, hPDE; melanocyte cells, hMEL) were treated with either dimethyl sulfoxide (DMSO, vehicle control) or varying concentrations of reversine (75 nM and 150 nM). Cells were analyzed at Day 2 for immediate chromosomal instability and selection and monitored through Day 28 to evaluate survival bottlenecks and proliferative competition. Outcomes were measured using KaryoTapR to capture chromosome-level and arm-level gains and losses. **(B-D)** Chromosome-level and arm-level instability induced by reversine treatment in mammary epithelial cells (hMEC, B), pancreatic epithelial cells (hPDE, C), and melanocyte cells (hMEL, D) at Day 2 and Day 28. Violin plots show the counts of chromosome and arm-level gains and losses across DMSO-treated and reversine-treated conditions (75 nM and 150 nM). Green represents Day 2, pink represents Day 28, indicating how chromosomal instability evolves. Statistical tests comparing condition groups are indicated (*p < 0.05, **p < 0.01, ****p < 0.0001; n.s.: not significant). **(E-J)** Correlation between chromosome size and aneuploidy frequency (gains and losses) following reversine treatment. Scatter plots display the normalized (reversine frequencies over DMSO control) aneuploidy frequency versus chromosome size for hMEC (E, H), hPDE (Panels F, I), and hMEL (G, J). E-G depict analyses at Day 2, and H-J illustrate Day 28. Regression lines represent trends for 75 nM and 150 nM reversine treatments; correlation coefficients (Pearson’s R) and p-values are provided for statistical significance.

Three near-diploid hTERT immortalized cell lines - hPDE (Human Pancreatic Duct Epithelial), hMEL (Human Melanocyte), and hMEC (Human Mammary Epithelial) - were used for this screen upon labeling them with a unique DNA barcode enabling demultiplexing (**Fig S8**). These cell lines were subjected to treatment with DMSO (control) or two concentrations of Reversine (75 nM and 150 nM) for up to 48 hours. Following Reversine treatment, cells were allowed to recover, and on the second day post-release (hereafter referred to as Day 2), a subset of cells was harvested and prepared for sequencing. This early timepoint provides an initial snapshot of the chromosomal landscape shortly after drug exposure. Cells were then cultured under standard conditions for an extended period of approximately 28 days, allowing evolutionary dynamics to unfold (**Fig 4A**). This later timepoint will be referred to as Day 28.

On Day 2, both 75 nM and 150 nM Reversine concentrations resulted in high levels of chromosome- and chromosome-arm gains and losses compared to the DMSO control, indicating that aneuploidy events are initially increased in response to Reversine treatment (**Fig 4B-D**). The total aneuploidy frequency (gains and losses) appeared to disproportionately involve larger chromosomes at Day 2, as suggested by significant positive correlations between normalized aneuploidy frequencies and chromosome size across all tested lines (**Fig 4E-G**). This indicates that chromosome size is an important factor in determining the likelihood that a chromosome undergoes missegregation. Notably, despite the observed positive correlations with chromosome size, we did not observe a similar trend for centromere size (**Fig S9**), indicating that shortly after the induction of missegregation through Reversine (Day 2), chromosome size rather than centromere size drives the bias toward chromosomal gains and losses. However, by Day 28, this correlation was notably reduced or absent in all tested cases, coupled with an overall decrease in aneuploidy levels in both treatment groups across all tested cell lines (**Fig 4H-J**). At Day 28, a significant reduction in the average number of gains and losses per cell was observed compared to Day 2 (**Fig 4B-D**). These findings underscore the presence of strong negative selective pressures acting against aneuploidy during prolonged *invitro* culture, favoring the restoration of genomic stability.

### Estimation of positive and negative selection on chromosomal gains and losses during cell proliferation

We aimed to utilize the frequencies of chromosome arm gains and losses at Day 2 and Day 28 to estimate whether their events were under positive or negative selection during in vitro proliferation. All analyses were performed at the chromosome-arm level, which represents the highest resolution achievable with KaryoTap and allows detection of both whole-chromosome and arm-level events. To quantify the magnitude of selection acting on gains and losses in each cell line, we computed chromosome arm selection scores. For each arm and event class (gain or loss), we performed a Fisher’s exact test to compare the Day 28 to Day 2 frequencies to derive a selection score. Selection scores were defined as the log2 odds ratio: positive (gain or loss) selection scores indicate enrichment of the event at Day 28 relative to Day 2 (positive selection), whereas negative values indicate depletion of the event at Day 28 relative to Day 2 (negative selection) (**Fig 5A-C, S10**, see also **Methods**). This framework accounts for both cell line-specific, pre-existing aneuploidies and chromosome-size-dependent missegregation frequencies, isolating and quantifying selection during the outgrowth phase.

**Figure 5.**
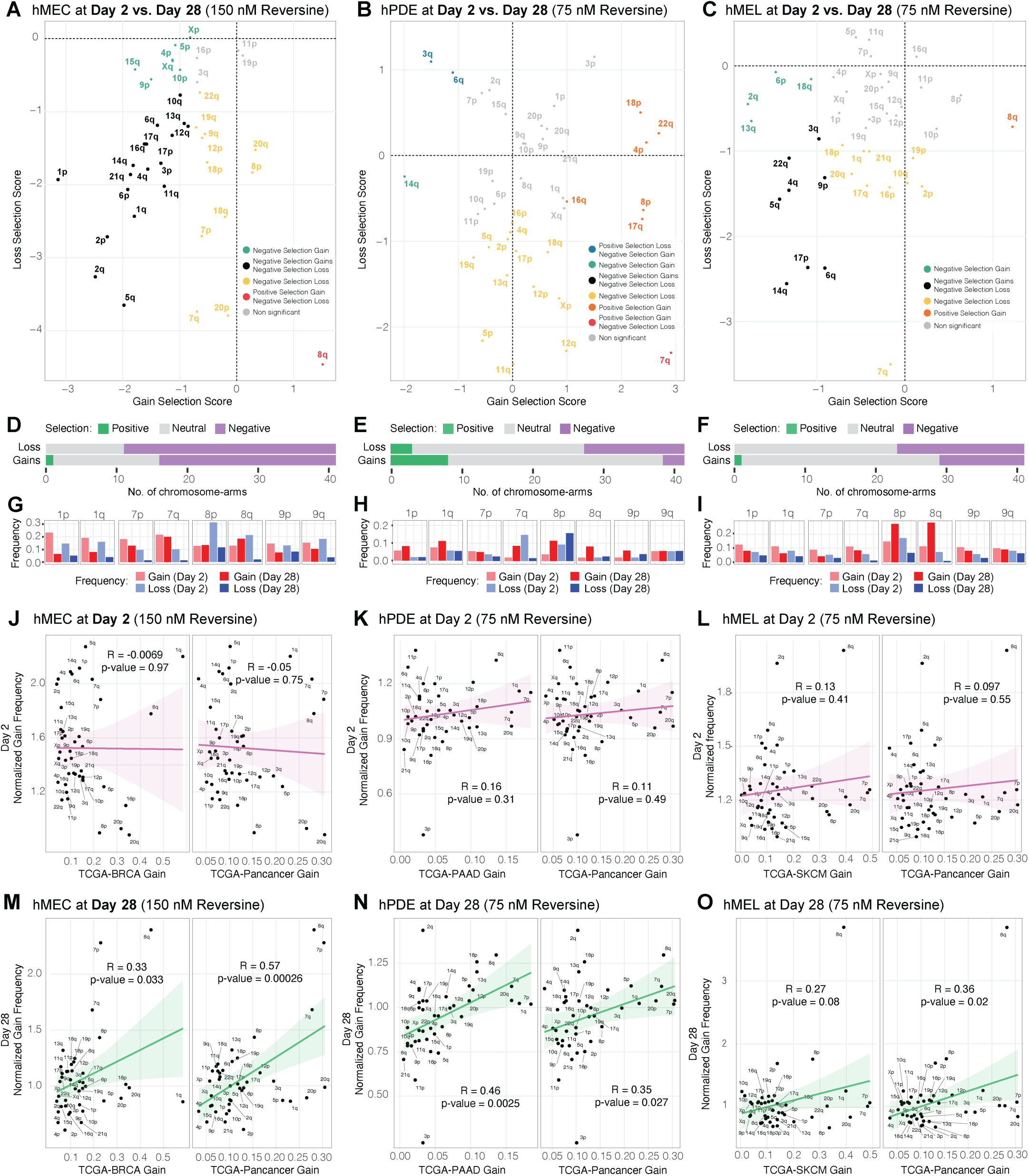
Quantification of selection forces and correlations between reversine-induced chromosomal alterations in cell lines and TCGA cancer data. **(A-C)** Selection scores for chromosome gains and losses across experimental conditions. Scatter plots display gain selection scores (x-axis) versus loss selection scores (y-axis) for hMEC (G, 150 nM reversine), hPDE (H, 75 nM reversine), and hMEL (I, 75 nM reversine). Positive selection (colored dots) indicates chromosomes that are preferentially gained (red or yellow) or lost (blue or turquoise) over time. Negative selection indicates chromosomes that are under negative selective pressure. Non-significant changes are indicated in gray. **(D-F)** Bar plot showing the number of chromosome arms under different types of selection (positive, neutral or negative) for arm gains and losses. **(G-I)** Frequency of gains and losses across specific chromosomal arms (of chromosome 1, 7, 8, and 9), at the two time points, Day 2 and Day 28. **(J-O)** Comparison of chromosome-arm gain frequencies induced by reversine with cancer amplification frequency from TCGA data. Panels compare normalized chromosome arm gain frequencies at Day 2 (A-C) and Day 28 (D-F) with TCGA amplifications from respective cancer types (hMEC with BRCA, hPDE with PAAD, and hMEL with SKCM) and TCGA-Pancancer. Scatter plots include correlation coefficients (Pearson’s R) and p-values. Green and pink regression lines distinguish between gains at Day 2 (pink) and Day 28 (green).

Overall, the majority of chromosome-arm copy-number events were classified as negatively selected across all conditions (**Fig 5A-I**). For example, the frequency of chromosome 8q loss decreased significantly from Day 2 to Day 28 in hMECs (from 21% at Day 2 to 2% at Day 28, **Fig 5G**); similarly chromosome 1q gain decreased from Day 2 to Day 28 in hMECs (from 19% at Day 2 to 7% at Day 28, **Fig 5G**). Despite widespread negative selection, a few events were also classified as positively selected (80 under negative and 12 positive across all lines), such as 8q gain in hMEL and hMEC, (respectively from 11% and 12% at Day 2 to 28% and 18% at Day 28, Fig. 5G, I); or 7q gain in hPDE (from 2% at Day 2 to 9% at Day 28, **Fig. 5H**). Overall, events under negative selection were dominated by losses (∼60% of negatively selected events were losses) while events under positive selection were for the vast majority gains (∼80% of them). In line with this asymmetry, losses exhibited a significantly lower frequency at Day 28 than Day 2 (**Fig 5G-I** and **Fig S10**).

Across the three epithelial lineages, we observed both common and cell type-specific selection landscapes. In hMEC and hMEL, but not hPDE, (**Fig 5A**), 8q gain was under strong positive selection. In addition, in hMEC there was significant negative selection for tissue-specific chromosome arm gains, such as 9p, while most of the other chromosome arms showed negative selection for both gains and losses. In hPDE (**Fig 5B**), 7q shows strong positive selection for gains and negative selection for losses, while 3q and 6q show strong negative selection for gains and positive selection for losses. hMELs (**Fig 5C**) show significant positive selection for gains in 8q and negative selection for gains in 2q, 6p, 13q, and 18q, and strong negative selection against losses in 1q, 7q, 17q, and 19p.

One notable example of common positive selection across multiple cell lines is 8q gain in both hMEC and hMEL cells, which showed a significant increase in frequency from Day 2 to Day 28 compared to the diploid state, suggesting that this event confers a proliferative advantage in both these cell types. Knowing the frequency at Day 2 and Day 28, along with their doubling times, we were able to estimate the fitness advantage conferred by 8q gain in these cell lines, deriving a selection coefficient of 0.0624 for hMEL and 0.0133 for hMEC (see **Methods**). These selection coefficients demonstrate tissue-specific fitness advantages conferred by 8q gain during in vitro proliferation. The nearly five-fold stronger selection in hMEL cells compared to hMEC cells indicates that identical chromosomal alterations can provide markedly different proliferative advantages depending on cellular context. While 8q is recurrently gained in both melanoma and breast cancer, our quantitative measurements suggest that its fitness contribution varies substantially across tissue types, potentially reflecting differences in the baseline expression or functional importance of 8q-resident genes like MYC in different epithelial lineages. Altogether, these analyses indicate that most aneuploidies – both gains and losses – are subject to negative selection during in vitro proliferation, with notable exceptions such as 8q gain and 7q gain.

### KaryoTap reveals that selection shape the patterns of gains observed in human cancer patient, with a more prominent role for positive selection

To investigate whether chromosomal evolution under experimental in vitro conditions reflects aneuploidy patterns observed in human cancers, we compared the chromosome-arm gain frequencies from TCGA (BRCA, PAAD, SKCM, and Pan-cancer) to the normalized Reversine-treatment gain frequencies (defined as the frequency in Reversine-treated cells over DMSO control), at both Day 2 and Day 28. Specifically, we asked whether the chromosome-arm aneuploidy observed in hMEC, hPDE, and hMEL cells over time would recapitulate the gain patterns characteristic of breast cancer (BRCA), pancreatic adenocarcinoma (PAAD), and skin cutaneous melanoma (SKCM), respectively, as well as in a pan-cancer analysis. The scatterplots in **Fig 5J-O** show the comparison between experimental gain frequencies and the corresponding TCGA gain frequency for each tissue at Day 2 (J-L) and Day 28 (M-O).

At Day 2, there was no significant correlation between the chromosome-arm gain frequencies in TCGA samples (BRCA, PAAD, SKCM, and Pan-cancer) and those observed in the matched *in vitro* counterparts (hMEC, hPDE, and hMEL). This indicates that the immediate effects of Reversine induce aneuploidy in a manner that does not mirror cancer-associated patterns, instead reflecting intrinsic missegregation propensities of specific chromosomes (**Fig 4E-J**). By Day 28, however, we observed a partial recapitulation of cancer-associated gain patterns: correlations between TCGA gain frequencies and the corresponding *in vitro* gain frequencies significantly increased across tissues. For example, the correlation between BRCA gain frequency and hMEC increased from −0.0069 at Day 2 (n.s.) to 0.33 at Day 28 (p < 0.05). Similarly, hPDE increased from 0.16 (n.s.) at Day 2 to 0.46 (p < 0.01) at Day 28, and hMEL from 0.13 (n.s.) at Day 2 to 0.27 (p = 0.08). This underscores the role of selection pressures that emerge over time during culture, progressively shaping tissue-specific aneuploidy landscapes toward cancer-like patterns. Notably, even at Day 28 the correlations, while significant, were moderate, indicating that *in vitro* evolution only partially recapitulates the aneuploidy landscape found in cancers. Moreover, these analyses focus on gains: we did not observe a similar recapitulation for losses (**Fig S11**; see **Discussion**).

Finally, we asked whether it was primarily the events under positive or negative selection that determined the significant correlation between the *in vitro* proliferation screen and the frequencies observed in TCGA. To address this, we excluded the events that were identified as being under positive or negative selection from the TCGA correlation analysis, as described above. We observed that when excluding events under negative selection, the significant association between the frequency of gains *in vitro* and in TCGA (Pancancer frequencies) remained intact across all tested cell lines (hMEC Pearson’s R from 0.33 to 0.51; hPDE Pearson’s R from 0.46 to 0.42; hMEL Pearson’s R from 0.27 to 0.25). However, strikingly, when excluding events under positive selection, the positive association between the *in vitro* screen and TCGA frequencies strongly decreased for all tested cell lines (hMEC Pearson’s R from 0.33 to 0.064; hPDE Pearson’s R from 0.46 to 0.32; hMEL Pearson’s R from 0.27 to -0.042). **Overall, this analysis suggests that positive selective pressure is a critical driving force shaping the patterns of chromosomal gains observed in human tumors.**

## Discussion

We designed custom panels for the Tapestri platform that enable targeted scDNA-seq of specific loci across all human chromosomes, coupled with a GMM-based copy number calling analysis pipeline for the identification of chromosome- and chromosome-arm-scale aneuploidy in thousands of cells with high accuracy and at greatly reduced sequencing depth compared to single-cell WGS methods(17–19). To increase ease-of-use, we compiled the computational scripts into an R package, *karyotapR*, which facilitates copy number calling and visualization (**Fig S12**). Using KaryoTap, we conducted in vitro proliferation screens that revealed how selection forces shape cancer genomes, demonstrating that proliferative selective pressures explain chromosomal gain patterns observed in tumors, while losses appear driven by non-proliferative cancer phenotypes. This easily adoptable, flexible, cost-efficient and highly scalable system lays the groundwork for an innovative class of tools for studying aneuploidy and chromosomal instability in healthy and diseased tissues and tumors.

While single-cell aneuploidy detection is not unique to KaryoTap, our design has several advantages, the most critical being the ability to scale up throughput to thousands of cells in one experimental run, thus significantly reducing hands-on time, reagent cost, and sequencing cost per cell. Additionally, targeting of specific loci allows us to forgo the typical technical difficulties of conventional WGS analysis, including correcting for mappability bias and GC bias, and the use of segmentation algorithms to make copy number calls(26). Furthermore, the total number of sequencing reads needed to obtain high-accuracy copy number calls for KaryoTap is greatly reduced compared to WGS-based scDNA-seq methods, and the targeted nature of the assay spreads the reads more evenly across the genome, preventing the biasing of detection sensitivity toward larger chromosomes that is seen with WGS. We expect this will be particularly important for accurately assessing chromosomal instability as the smallest of chromosomes will be more equally represented in the data relative to the larger chromosomes. Finally, the commercial availability of the Tapestri system allows for easier adoption and reproducibility compared to non-commercialized "homebrew" methods that need to be established and optimized in each lab from scratch (22).

As evidenced by the analysis of RPE1, our method generates a range of smooth copy number measurements for chromosomes with the same copy number, indicating some level of technical error. To account for this error and convert the continuous smooth copy number scores to discrete copy number values, we used a Gaussian mixture model (GMM) classification strategy, which has been previously used for copy number analysis of single-cell whole genome sequencing data(34). This allows each smooth copy number score to be associated with a set of (posterior) probabilities of being measured from a chromosome of a given range of copy numbers (e.g., 1, 2, 3, 4, or 5). As the number of probes increases, the variance of the model components decreases, resulting in an increase in classifier accuracy that we observe between our iterative v1 to v3 panel designs. While we use a near-diploid cell line as our ground truth, the GMM strategy also allows us to calculate the expected (theoretical) sensitivity that a chromosome with copy number 1, 2, 3, 4, or 5 would be correctly called using our system by calculating the proportion of overlap between the copy number components of the model. Further improving the panel by increasing the number of probes will allow us to more confidently observe the karyotype heterogeneity in these samples as well as in tumors and other tissues.

Deep sequencing (∼80-100X on average per cell) of each target region allows for robust single nucleotide variant (SNV) calling that is not possible at the lower genomic coverage afforded by other high-throughput methods(22). This enabled us to resolve and identify 5 multiplexed cell lines in a single experiment by clustering cells by variant allele frequencies. Since our panels specifically target loci known to harbor SNPs across the human population, we can extend sample multiplexing to clinical samples (e.g., tumor tissue) from different individuals without the need for barcodes, further reducing experimental costs. Additionally, while our panels were designed for copy number analysis, additional probes could be added that cover tumor suppressor genes and oncogenes of interest, thus revealing consequential point mutations alongside chromosomal copy number. Mission Bio offers several ready-made panels covering mutational hotspots and genes relevant to a broad range of tumor types, allowing for a great degree of customizability. Furthermore, we demonstrated that SNV-associated allele frequency shifts detected using KaryoTap could be used to infer loss of heterozygosity (LOH), a common event in cancer where the germline heterozygous state of a chromosome changes to a homozygous state in tumor cells(33). LOH has been demonstrated to promote tumorigenesis by inactivating tumor suppressor genes through chromosomal loss(31,32). In the context of chromosomal instability, the chromosome remaining after the loss of its homologue can be duplicated, resulting in LOH with a net-neutral copy number change (copy number neutral (CNN)-LOH). The deep sequencing depth and copy number detection enabled by KaryoTap allow for discovery of CNN-LOH events, which would otherwise be difficult to detect with the shallow coverage typical of other scDNA-seq methods(33).

To enable experimental design flexibility when using our custom panels, we added a set of probes that can detect DNA barcode and CRISPR gRNA sequences integrated into the genome. DNA barcodes can be used to multiplex and resolve cells belonging to different samples in a single Tapestri experiment that otherwise could not be distinguished by genotype. Through barcoding, users can compare samples from the same individual or compare experimental and control conditions in the same cell lines while minimizing batch effects. Regardless of design, combining several samples into one experimental run greatly reduces the per-sample reagent and sequencing costs in addition to the hands-on time required to process the samples. Detecting barcodes in thousands of cells is made possible by exploiting the targeted nature of the sequencing assay. Single-cell DNA sequencing methods with comparable throughput rely on inefficient and random transposon insertion, which would only detect a randomly inserted barcode in about 20% of cells(22). By specifically targeting and selecting for the barcoded insert, we can reliably recover the barcode sequence in over 90% of cells. Furthermore, including a probe that targets inserted CRISPR gRNA sequences allows for an additional layer of experimental design flexibility where the gRNA-mediated treatment that each cell receives can be identified by the gRNA sequence itself. Here, we demonstrated the gRNA detection and multiplexing capabilities of our system by evaluating the efficiency and specificity of KaryoCreate, our method for inducing chromosome specific aneuploidy. Since the gRNAs can be detected in 70-80% of cells when transduced at low MOI, this system could also be used for CRISPR screen applications where cells are randomly treated with one gRNA from a library of hundreds of possible gRNAs and thus require high detection sensitivity(35). Altogether we describe KaryoTap as a cost-effective, high-throughput single-cell DNA sequencing method to detect chromosome-scale aneuploidy with high accuracy and enable functional genomic screens through barcode/gRNA detection.

A fundamental question in understanding chromosomal instability is what structural parameters of chromosomes determine their propensity for missegregation. Our data reveal a strong positive correlation between chromosome size and missegregation frequency at Day 2 following Reversine treatment, with larger chromosomes exhibiting higher rates of gains and losses across all three cell lines tested (**Fig 4E-G**). Chromosome size is often associated with nuclear positioning, as larger chromosomes tend to occupy more peripheral locations within the nucleus. A recent study demonstrated that nuclear chromosome location dictates segregation error frequencies, with chromosomes in certain nuclear positions being more prone to entrapment in micronuclei following error-prone mitosis (69). Our findings are consistent with a model in which chromosome size—and its associated nuclear positioning—influences the likelihood of missegregation during mitosis when missegregation is induced via MPS1 inhibitors. Notably, we did not observe an association between centromere size and missegregation frequency (**Fig S7**) (65). This discrepancy may reflect differences in the mechanisms of missegregation operating in different cell and tissue types. Interestingly, gain of 1q, one of the most frequent events observed in our study (Figure 5), is also frequently found in the normal mammary gland; thus its presence in normal mammary tissue may reflect both the frequency of occurrence and a potential selective advantage (70, 75).

Our study reveals that the majority of aneuploidies—both gains and losses—are subject to negative selection during *in vitro* proliferation. While positive selection enriches specific chromosomal gains such as 8q and 20q, the broader aneuploidy landscape is dominated by the elimination of deleterious aneuploidies over time. By comparing our in vitro screen with primary tumor data, we observed that the frequency of chromosomal gains at Day 28 (but not Day 2) significantly correlated with gain frequencies in matched TCGA cancer types. Most importantly, when we excluded chromosomal events identified as being under positive selection, these correlations decreased or disappeared, whereas excluding negatively selected events mostly preserved the TCGA associations. This demonstrates that the patterns of chromosomal gains observed in human cancers are shaped predominantly by selective pressures acting during proliferation, rather than by the intrinsic frequency of chromosome missegregation. The convergence between our *in vitro* evolution screens and patient cancer genome landscapes thus reflects shared selective advantages operating in both contexts. Collectively, these findings demonstrate that proliferative selection shapes the landscape of chromosomal gains in cancer, with positive selection exerting a more prominent influence than negative selection in determining the final aneuploidy patterns. Interestingly, in a complementary study (77), computational inference of positive and negative selection across cancer aneuploidy revealed that positive selection scores are positively associated with Day 28 gain frequencies, while negative selection scores are negatively associated with Day 28 gain frequencies. In agreement with our findings, positive selection scores showed stronger correlations than negative selection scores (77).

The chromosome 8q and 20q gains we observed under positive selection in hMEC cells align with well-established prognostic markers in breast cancer associated with recurrence and poor clinical outcomes (63, 66). Similarly, the frequent selection of chromosome 20 gain in hPDE cells mirrors the recurrent amplification of this chromosome in pancreatic adenocarcinoma (62). These findings suggest that both positive and negative selection cooperate to sculpt the tissue-specific aneuploidy landscapes characteristic of different cancer types.

Our chromosomal instability screens also converge with recent work (72) in demonstrating tissue-specific selection of chromosome 8q gain in mammary epithelial cells, with our quantitative fitness measurements (s = 0.0133 in hMEC, s = 0.0624 in hMEL) supporting the proposed MYC-driven proliferative advantage. However, unlike this prior study, we did not observe positive selection for chromosome 1q gain; in fact, +1q showed negative selection in our bulk competition assays. This divergence likely reflects fundamental differences in competitive dynamics between clonal isolation (72) and mixed-population selection (this study). Additionally, the 21 population doublings in our screen may be insufficient for the Notch poising phenotype to overcome the initial fitness costs of aneuploidy, whereas the 35-40 PD timeframe and pre-established aneuploid backgrounds in the Watson et al. study may allow stress-mitigation effects to emerge. This suggests that +1q provides a selective advantage specifically in competitive scenarios where +1q-bearing cells remain rare within a population—a condition more likely to occur in spatially structured tissues or during clonal evolution from single transformed cells than in bulk culture competition.

A striking observation from our study is the asymmetry between chromosomal gains and losses: while in vitro gain frequencies at Day 28 correlated significantly with TCGA tumor profiles, no such association was observed for losses. This suggests that the selective pressures driving chromosomal losses in tumors differ fundamentally from those acting on gains. We propose two non-mutually exclusive explanations for this discordance. First, chromosomal losses may impose stronger deleterious effects on cell viability in vitro, requiring additional oncogenic events—such as TP53 inactivation or whole-genome doubling—to be tolerated during two-dimensional culture. Second, chromosomal losses may be selected primarily by in vivo-specific pressures such as immune evasion, metabolic reprogramming, or metastatic capacity, rather than by the proliferative advantage that drives gains in standard culture conditions (76). Consistent with this interpretation, recent studies have documented the accumulation of cells harboring specific chromosomal losses—including 10q, 16q, and 22q—in normal aging tissues such as the mammary gland (70, 75). Notably, these loss events were not observed in our in vitro screens or in other long-term two-dimensional evolution experiments (72), suggesting they are tolerated within three-dimensional tissue architecture but do not confer proliferative advantages under standard culture conditions. Supporting the importance of tissue context, gastric epithelial cells cultured as three-dimensional organoids (with TP53 inactivation) frequently acquired 3p loss, a hallmark alteration in gastric cancer (78). Together, these observations suggest that the tissue microenvironment—including stromal interactions, immune surveillance, and spatial constraints—plays a critical role in determining which aneuploidies are ultimately selected during tumor evolution in vivo.

## Methods

### Cell Culture

All cells were grown at 37°C with 5% CO2 levels. All cell media was supplemented with 1X pen-strep, and 1X L-glutamine. hTERT human retinal pigment epithelial cells (RPE-1; ATCC CRL-4000) and SW48 cells (ATCC CCL-231) were incubated in DMEM, supplemented with 10% FBS. LoVo cells (ATCC CCL-229) were incubated in Ham’s F12-K media with 10% FBS. LS513 cells (ATCC CRL-2134) and hTERT human pancreatic nestin-expressing cells (hPNEs; ATCC CRL-4023) were incubated in RPMI media with 10% FBS. CL11 cells (Cellosaurus CVCL_1978) were incubated in DMEM:F12 and 20% FBS. hTERT p53-/- human colonic epithelial cells (hCECs; Ly et al.(36)) were cultured in a 4:1 mix of DMEM:Medium 199, supplemented with 2% FBS, 5 ng/mL EGF, 1 μg/mL hydrocortisone, 10 μg/mL insulin, 2 μg/mL transferrin, 5 nM sodium selenite, pen-strep, and L-glutamine. hTERT human mammary epithelial cells (hMEC) (67) were incubated in Mammary Epithelial Cell Growth Medium (MEGM, Lonza) supplemented with 1% FBS and the MEGM SingleQuots kit (Lonza). hTERT human pancreatic duct epithelial cells (hPDE; Cellosaurus CVCL_4376) were cultured with Gibco keratinocyte serum-free medium with 10% FBS, 25 mg of bovine pituitary extract, and 2.5 μg of human recombinant EGF. Human melanocytes (hMEL) with hTERT overexpression and p16/CDKN2A knocked out (gRNA: CCCAACGCACCGAATAGTTACGG) were obtained from Richard Marais lab (University of Manchester) were cultured with Melanocyte Growth Medium (MGM, Cell Applications) supplemented with 1% FBS. For long-term storage, cells were cryopreserved at −80°C in 70% cell medium, 20% FBS, and 10% DMSO. All cell lines were tested for mycoplasma.

### Custom Tapestri Panel Design

Panel Version 1 (CO261) comprises 330 probes across the 22 human autosomes and the X chromosome. To identify candidate target regions for the panel, we used the Common SNP files downloaded from UCSC(37,38) (snp151Common, hg19), and considered only synonymous variant SNPs with a major allele frequency at >0.5 and <0.6. For cytobands with more than 4 synonymous variants, we split the cytoband into 4 subregions based on the percentile of the cytoband coordinates (0-25th percentile, 25-50th percentile, 50-75th percentile, and 75-100th percentile). From each subregion, we randomly selected 1 SNP as a representative candidate. In cases where there were less than 5 synonymous variant SNPs, all SNPs were used. We submitted all candidate SNPs to the Tapestri Panel Designer to generate a panel design and ensured that the designed probes targeted the candidate SNPs and had similar GC contents. Next, randomly selected probes such that each chromosome had a probe density of ∼1 per 10MB. Panel Version 2 (CO610) comprises 352 probes across all 24 human chromosomes. This panel was generated using Panel v1 as a base: first, we removed 61 probes that had low PCR amplification efficiency based on total read counts per probe. Then we added 82 probes such that each chromosome was targeted by at least 12 probes and included 4 probes targeting chrY. To enable detection of lentiviral-delivered gRNAs, we added one probe targeting the region of the construct containing the gRNA sequence and one probe targeting a region upstream as a vector control. Similarly, for detection of lentiviral-delivered DNA barcodes, we added one probe targeting the region of the construct surrounding the barcode sequence, and one probe targeting a region downstream as a vector control. Support for the custom panel design and synthesis of the panel was provided by Mission Bio (San Francisco, CA, USA). Panel maps were created using the karyoploteR R package(39).

### Tapestri Single Cell DNA Sequencing

Cell lines were trypsinized for 2-3 minutes, washed in room temperature Mg^2+^/Ca^2+^-free DPBS, centrifuged at 300g for 5 minutes, and resuspended in DPBS at a concentration of 3K cells/uL. For the experiment using the RPE1, SW48, LS513, LoVo, and CL11 cell lines, 600K cells from each cell line were combined, centrifuged at 300g for 5 minutes, and resuspended in Tapestri Cell Buffer at a concentration of 3.5K cells/uL. For the experiment using the RPE1, hPNE, and hCEC cell lines, 45K cells from each cell line were combined, centrifuged at 300 x g for 5 minutes, and resuspended in Tapestri Cell Buffer at a concentration of 4K cells/uL. For the KaryoCreate experiment, ∼100K cells from each condition were combined, centrifuged at 300 x g for 5 minutes, and resuspended in Tapestri Cell Buffer at a concentration of 3.4K cells/uL. For the Aneuploidy induction experiment, ∼200K cells from each condition were combined, centrifuged at 300 x g for 5 minutes and resuspended in Tapestri Cell Buffer at a concentration of 5K cell/ul. Cell droplet encapsulation, barcoding, and sequencing library preparation were performed using the Tapestri instrument according to the manufacturer’s instructions (Mission Bio, San Francisco, CA, USA). Sequencing was performed using an Illumina NovaSeq 6000 or NextSeq 500 in 2x150bp paired-end format. After sequencing, deconvolution of barcodes, read counting, and variant calling were handled by the online Tapestri Pipeline (v2.0.2)(25). The pipeline outputs both read counts per probe for each cell and variant allele frequencies for called variants for each cell.

### Low Pass Whole Genome Sequencing & Karyotyping

Genomic DNA was extracted from cell pellets using 0.3 μg/μL Proteinase K (QIAGEN #19131) in 10mM Tris pH 8.0 for 1 hour at 55°C, following heat inactivation at 70°C for 10 minutes. DNA was digested using NEBNext dsDNA Fragmentase (NEB #M0348S) for 25 minutes at 37°C followed by magnetic DNA bead cleanup with 2X Sera-Mag Select Beads (Cytiva #29343045). Library prep was performed using NEBNext Ultra II DNA Library Prep Kit for Illumina (NEB #E7103) according to the manufacturer’s instructions, generating DNA libraries with an average library size of 320 bp. Quantification was performed using a Qubit 2.0 fluorometer (Invitrogen #Q32866) and the Qubit dsDNA HS kit (Invitrogen #Q32854). Libraries were sequenced on an Illumina NextSeq 500 at a target depth of 4-8 million reads. Reads were trimmed using trimmomatic(40), aligned to the hg38 genome using bwa-mem(41), and analyzed for copy number variants using the CopywriteR(42) R package. G-banded karyotyping of 100 RPE-1 cells was performed by WiCell Research Institute, Inc. (Madison, WI).

### Cloning of sgRNAs

We modified the scaffold sequence of pLentiGuide-Puro (Addgene #52963) by Gibson assembly to contain the A-U flip (F) and hairpin extension (E) described by Chen et al(43). for improved sgRNA-dCas9 assembly, obtaining pLentiGuide-Puro-FE. sgRNAs were designed and cloned into this pLentiGuide-Puro-FE vector according to the Zhang Lab General Cloning Protocol(44). To be suitable for cloning into *BbsI*-digested vectors, sense oligos were designed with a CACC 5’ overhang and antisense oligos were designed with an AAAC 5’ overhang. The sense and antisense oligos were annealed, phosphorylated, and ligated into *BbsI*-digested pLentiGuide-Puro-FE for KaryoCreate purposes. Sequences were confirmed by Sanger sequencing.

### Gateway Recombination Cloning for Generation of Barcode Library

pHAGE-CMV-DEST-PGKpuro-C-BC was a library of lentiviral vectors containing 24-bp random barcodes that was built as described in Sack & Davoli et al., 2018(45). Destination vector pHAGE-CMV-DEST-PGKpuro-C-BC and entry vector pDONR223_BFP (Addgene: 25891) are recombined following the manufacturer’s protocol. Briefly, 50 ng of entry vector and 100 ng of destination vector are mixed with LR Clonase™ enzyme and incubated overnight at room temperature. The next day, the reaction mixture is incubated with Proteinase K at 37°C for 10 minutes, followed by inactivation at 75°C for 15 minutes. The reaction is then transformed into stbl3 bacterial competent cells, plated onto LB agar plates, and incubated overnight at 37°C. Individual clones are collected into 96 well plates and expanded. Plasmid is extracted from the bacterial culture using a 96-well mini-prep kit (Zymo kit, Zyppy 96 plasmid kit). All clones were sequenced by Sanger sequencing at the site of the barcode using primer ACTTGTGTAGCGCCAAGTGC. Duplicates were eliminated, and unique barcodes were retained in the final library. BFP expression and Puromycin selection were validated by transfecting randomly selected clones into HEK293T cells.

### Lentivirus Production and Nucleofection

For transduction of cells, lentivirus was generated as follows: 1 million 293T cells were seeded in a 6-well plate 24 hours before transfection. The cells were transfected with a mixture of gene transfer plasmid (2 μg) and packaging plasmids including 0.6 μg ENV (VSV-G; addgene #8454), 1 μg Packaging (pMDLg/pRRE; addgene #12251), and 0.5 μg pRSV-REV (addgene #12253) along with CaCl2 and 2× HBS or using Lipofectamine 3000 (Thermo #L3000075).The medium was changed 6 hours later and virus was collected 48 hours after transfection by filtering the medium through a 0.45-μm filter. Polybrene (1:1000) was added to the filtered medium before infection.

### KaryoCreate Experiments

KaryoCreate experiments were performed as described in Bosco et al., 2023(14). Briefly, p53-/- hCEC were first lentivirally transduced with pHAGE-KNL1Mut-dCas9 and selected with blasticidin. The cells were then lentivirally transduced with the indicated sgRNAs and selected with puromycin. scDNA seq was performed ∼10 days after transduction with the gRNAs. The sequences of the gRNAs targeting the centromeres of specific chromosomes are listed in Table S4 and were designed as described in Bosco et al., 2023(14). To compare conditions, Fisher’s exact test was performed in R using the ‘fisher.test()’ function, comparing the proportion of cells for each chromosome and sample that are diploid (copy number = 2) and aneuploid (copy number = {1, 3, or 4} between the sgNC control and sample and the given experiment sample. The Benjamini-Hochberg correction for multiple comparisons was applied to p-values using ‘p.adjust()’.

### Single Cell Aneuploidy Induction Screens and Selection Score Calculation

Human immortalized cell lines (hPDE, hMEC, hMEL) were lentivirally transduced with the demultiplexing barcode plasmids, as specified in Supplementary Table 5, and further selected with puromycin. A total of 1e+07 cells were treated with 1.5 μl of DMSO (control) or reversine (75nM or 150nM) for 36 to 48 hours. After reversine treatment, cells were split and allowed to recover, with medium changes every 2-3 days.

Cells were passaged at 80-90% confluence using Gibco TrypLE Express Enzyme (1X). Cells were frozen 48 hours and 28 days after reversine release (Day 2 and Day 28, respectively) in freezing medium containing 10% DMSO and 20% FBS in the respective growth medium.

To calculate the selection scores (https://github.com/fabio-alfieri/copy-number-annotation) for each arm, a 2×2 contingency table was constructed, comparing the frequency of gain (or loss) events between two experimental conditions (e.g., Day 2 vs. Day 28 at 150 nM Reversine). Fisher’s exact tests were then applied separately for gains and losses to assess whether the frequency of each alteration significantly changed between time points. P-values were adjusted for multiple testing using the Benjamini–Hochberg procedure. To quantify the direction and magnitude of selection, we computed selection scores as the logarithm of Fisher’s test odds ratios for gain and loss events, respectively. Positive scores indicate selection for a specific copy-number alteration (gain or loss) over time, whereas negative scores denote counter-selection.

### Calculation of Fitness Advantage from Aneuploidy Frequency Changes

To estimate the selective advantage conferred by specific chromosomal aneuploidies during in vitro culture, we calculated selection coefficients (s) from the change in aneuploidy frequency between Day 2 and Day 28 following reversine treatment. The selection coefficient represents the relative fitness advantage per cell generation, where a fitness of (1 + s) indicates the proportional increase in reproductive success compared to diploid cells.

We assumed a cell doubling time of 30 hours (1.25 days) based on growth curves for each cell line under standard culture conditions. Over the 26-day interval between Day 2 and Day 28, cells underwent approximately 20.8 cell divisions (26 days ÷ 1.25 days per division).

Under a model of constant selection, the frequency of an aneuploid population changes according to:

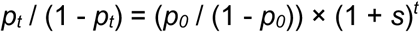

where *p0* is the initial frequency of the aneuploid population at Day 2, *pt* is the frequency at Day 28, *t* is the number of cell generations (20.8), and *s* is the selection coefficient.

For chromosome 8q gain in hMEC cells:

- Day 2 frequency (*p0*): 16% (0.16)
- Day 28 frequency (*pt*): 20% (0.20)
- Number of generations (*t*): 20.8 Solving for *s*: *s* ≈ 0.0133 (1.33% per generation)

A selection coefficient of *s* = 0.0133 indicates that chr8q gain cells have approximately 1.33% higher fitness per cell division compared to diploid cells under these culture conditions. Assuming that the fitness increase is due to decrease in doubling time, this would correspond to a 29.7 hours doubling time compared to 30.

For chromosome 8q gain in hMEL cells:

- Day 2 frequency (*p0*): 9% (0.09)
- Day 28 frequency (*pt*): 26% (0.26)
- Number of generations (*t*): 20.8

Solving for *s*: *s*≈0.0624 (6.24% per generation). In hMEL cells, the chr8q gain shows a substantially stronger selective advantage (*s*= 0.0624 or 6.24% per generation) compared to hMEC cells. Assuming that the fitness increase is due to decrease in doubling time, this would correspond to a 28.2 hours doubling time compared to 30.

### Parsing and Counting of Barcoded Reads

To detect specific gRNA or DNA barcode sequences, we searched for the known sequences against the cell-associated aligned reads (cells.bam file) generated from the Tapestri Pipeline. Search queries were conducted vcountPattern() using the Biostrings R package(46), with tolerance for up to 2 base mismatches. BAM files were manipulated using the Rsamtools R package(47).

### Cell Line Demultiplexing and Identification

To demultiplex cells from different cell lines in the initial scDNA-seq experiment, we use the allele frequency (AF) of variants that are called by GATC as part of the Tapestri Pipeline. Variants were filtered by selecting those with standard deviations of AF >20 to select variants whose allele frequencies vary the most across all cell lines. PCA was used to reduce the dimensions of the remaining variants. The top 4 principal components were embedded in two dimensions by UMAP and then clustered using the dbscan method. The 5 clusters with the greatest number of cells were kept, corresponding to the 5 expected cell lines. The remaining clusters, likely representing cell doublets, were discarded from further analyses. This method was repeated for subsequent Tapestri experiments, adjusting for the expected number of cell populations.

The cluster containing RPE1 cells in each experiment was identified by clustering with published deep WGS of RPE1. Published RPE1 WGS data was obtained from SRA Accession ERR7477340(28,48). Reads were aligned to the hg19 genome using bwa. MarkDuplicatesSpark and HaplotypeCaller from GATK were used to mark duplicate reads and get AFs from called variants(49). The vcfR R package(50) was used to extract the AFs for called variants common to the published data and our dataset. The mean AF for each variant was calculated for each of the 5 cell lines. PCA was used to cluster our mean AF dataset with the RPE1 AFs. The cell line that clustered most closely with the published RPE1 data was labeled as RPE1 cells.

### Copy Number Calling

Copy number scores for each probe in each cell (cell-probe scores) were calculated relative to RPE1, for which we know the copy number of each chromosome. The raw count matrix was normalized using two alternative approaches: (A) by scaling each cell’s mean to 1 (Equation 1) and then each probe’s median to 1 (Equation 2), or (B) by scaling each cell to cell’s total read count and then rescaling to a global median of 1 (Equation 3) and by correcting to the probe-specific variability where each probe was adjusted using a coefficient *cprobe* proportional to its dispersion across cells (Equation 4). The normalized counts were then scaled such that the value of the median normalized RPE1 counts for each probe was set to 2 for all probes except those targeting chr10q, which were set to 3 (Equation 5). The identities of the remaining 4 populations of cells were identified by comparing their overall copy number profile with matched bulk WGS data.

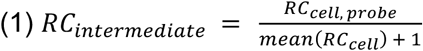

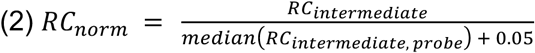

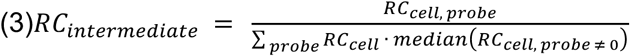

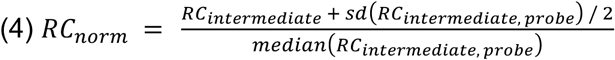

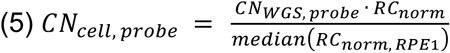

Smooth copy number scores for each chromosome in each cell (cell-chromosome scores) were generated by taking the median of the probe-specific copy number values for probes targeting a common chromosome (Equation 6). This was also modified to calculate cell-chromosome-arm scores for probes common to a chromosome arm

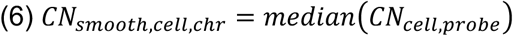

For all probes on chromosome *chr*.

Integer copy number values for each cell-chromosome were classified using Gaussian mixture models (GMMs) with either five components representing possible copy number values of 1, 2, 3, 4, and 5, or three components representing copy number values 1, 2, and 3. To generate the GMMs, the normalized counts for each probe for the RPE1 cells were fitted to Weibull distributions using the fitdistrplus R package(51). These Weibull parameters represented parameters for copy number = 2 for all probes except those targeting chr10q, which has 3 copies in RPE1. The scale parameters were then scaled for possible copy number values 1 through 6, relative to the RPE1 copy number: probes with RPE1 copy number = 2 were scaled by 50%, 100%, 150%, 200%, 250%, and 300% for copy number = 1, 2, 3, 4, 5 and 6; probes with RPE1 copy number = 3 were scaled by 33%, 67%, 100%, 133%, 167%, and 200% for copy number = 1, 2, 3, 4, 5 and 6. 500 Weibull-distributed values are drawn using each of the six parameter sets for each probe to simulate six matrices of 500 simulated cells. For each cell, the values were smoothed across the probes belonging to each chromosome to simulate cell-chromosome copy number values. The distribution of the scores for each chromosome was then fit to Gaussian (normal) distributions, separately for each copy number level. The result is a set of normal parameters (mean μ and standard deviation σ) for each chromosome for each value of copy number *k* = {1, 2, 3, 4, 5, 6}. The six copy number Gaussian components for each chromosome were combined into a GMM, representing the probability densities for each copy number value for that chromosome (Equation 7). Using Bayes rule and assuming equal priors, the posterior probability of a cell-chromosome copy number score being generated under each component *k* is given by Equation 8.

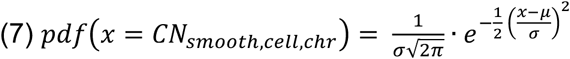

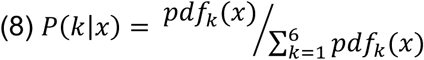

Decision boundaries for the GMMs are calculated by finding the transitions between components, i.e., the point x where the probability density functions (PDFs) of the components are equal. We evaluated copy numbers using GMMs including copy number components 1-5 throughout the study and 1-3 where indicated. Upper boundaries for component 5 were calculated using components 1-6. Theoretical sensitivity for each copy number component was calculated as the proportion of the component PDF that falls within its decision boundaries (i.e., true positive rate). 5-fold cross validation was performed by partitioning RPE1 cells into 5 equally sized groups and using each group once to evaluate a model generated using the remaining 4 groups. R scripts for copy number calling were compiled into an R package, karyotapR. karyotapR version 0.1 was used for analyses in this study.

### Panel Simulations

For the probe downsampling simulation of chromosome 2, 50 samples each of *n* probes from the set of 24 probes targeting chr2 were generated where *n* = {4, 6, 8 … 20, 22}. Copy numbers were called for each set of probes for each cell. The sensitivity of the copy number calls was recalculated for the RPE1 cells as well as the theoretical sensitivity for all GMM components. This analysis was repeated for the set of probes targeting chr6 where *n* = {4, 6, 8 … 14, 16}.

For the simulation of an expanded custom panel, the set of 120 probes targeting chromosomes 1, 2, 3, 4, 5, and 6 were sampled 50 times to produce 50 sets of the 120 probes in unique orders. Starting with the first 4 probes of each set, copy numbers were called for RPE1 cells using those 4 probes, and again as each additional probe was added to the set until all 120 probes were used for the calculation. This procedure was repeated for each of the 50 sets. The sensitivity of copy number cells was recalculated for the RPE1 cells at each step as well as the theoretical sensitivity for all GMM components using a model with 5 components (with the upper boundary of the 5th component being calculated using a 6-component model), and with 3 components where noted.

For the read depth simulation, the read count matrix was randomly downsampled to 80%, 60%, 40%, and 20% of the total read counts using the downsampleMatrix function from the scuttle R package(52).

50 downsampled matrices were generated for each downsampling level. The sensitivity of copy number cells was recalculated for the RPE1 cells at each step as well as the theoretical sensitivity for all GMM components using a 5-component model. Least squares linear regression was performed at each copy number level in R with the formula ’sensitivity ∼ log(n.probes) + depth’ where depth was a categorical variable.

### Comparison between KaryoTap and Arcwell

We compared the copy number detection ability of KaryoTap with Arc-well, a scDNA-seq method of similar throughput. Arc-well fastq files were aligned to the reference human genome hg38 using BWA-mem (v0.7.17), and duplicates were removed using GATK (Genome Analysis Toolkit, v4.1.7.0) to generate analysis-ready BAM files. To enable fair comparison with KaryoTap sequencing depth, Arc-well data were downsampled to 30K and 15K reads per cell. The resulting BAM files were processed using the R package CopywriteR (v1.18.0) to determine arm-level copy numbers. Downsampling to 15K reads resulted in qualitatively increased noisiness, with only 30-50% of full-depth copy number events detected. For comparison, we downsampled KaryoTap RPE1 data to 40X depth (∼15K reads per cell). For cost comparison, we evaluated reagent costs based on current pricing. GenomePlex WGA4, the DOP-PCR WGA product (60), costs $19.45–27.70 per cell, pricing a hypothetical 500-cell experiment at $9,725 before sequencing costs. In contrast, KaryoTap costs approximately $0.50–1.00 per cell, representing less than 10% of the cost of alternative methods such as shallow scWGS based on single-well cell capture, which costs approximately $8–10 per cell.

## Supplementary Data

### Systematic Evaluation of Technical Parameters Influencing Copy Number Classification Performance

To comprehensively characterize the factors governing copy number calling accuracy in KaryoTap, we conducted a series of systematic simulations using scDNA-seq data from RPE1 cells. These simulations examined three key technical parameters: probe number per chromosome, copy number classification scheme (5-component versus 3-component Gaussian mixture models), and sequencing depth. By independently varying each parameter while controlling for others, we aimed to quantify their relative contributions to classification sensitivity and guide optimization strategies for future panel designs.

### Probe Number Determines Classification Sensitivity

To confirm that copy number classification accuracy depends on the number of probes targeting a chromosome rather than the absolute size of the chromosome itself, we performed downsampling analysis on chromosomes 2 and 6. We randomly sampled n probes from the complete probe sets targeting chromosome 2 (24 total probes) and chromosome 6 (18 total probes), generating 50 independent samples for each value of n. Copy number classification accuracy was recalculated for RPE1 cells using each downsampled probe set.

The downsampling analysis revealed a clear relationship between probe number and classification performance. For chromosome 2, median accuracy decreased progressively from 95.8% with the full complement of 24 probes to 68.8% with only 4 probes. Similarly, chromosome 6 showed a decline from 91.4% (18 probes) to 66.8% (4 probes), confirming that probe number, not chromosome size, is the primary determinant of classification accuracy (**Fig S5A**).

Importantly, the variability of classification accuracy also increased markedly as probe number decreased. The interquartile range (IQR) of accuracy distributions expanded from 0.8 percentage points with 22 probes to 9.14 percentage points with 4 probes for chromosome 2, and from 1.5 percentage points with 16 probes to 11.6 percentage points with 4 probes for chromosome 6 (**Fig S5A**). This increasing variability indicates that sparse probe coverage not only reduces average accuracy but also introduces greater inconsistency in classification performance across independent samples.

These patterns of decreasing accuracy and increasing variance with fewer probes were consistently observed across all five copy number levels (1-5 copies) when examining theoretical sensitivity calculated from the Gaussian mixture model components (**Fig S5B**). This consistency across copy number states demonstrates that the relationship between probe number and classification performance is a fundamental property of the method rather than an artifact specific to diploid chromosomes.

### Simulated Panel Expansion Defines Optimal Probe Requirements

Having established that increasing probe number enhances sensitivity, we sought to determine the optimal probe coverage required to achieve high classification accuracy across all copy number states. To address this question, we simulated progressively larger panels by combining all probes targeting chromosomes 1 through 6 (120 probes total) and treating them as measurements from a single hypothetical chromosome. This approach is valid because all six chromosomes are diploid in RPE1 cells, allowing us to pool their probe measurements while maintaining a known ground truth of 2 copies.

For each of 50 simulation trials, we constructed panels by iteratively sampling the 120 probes without replacement, recalculating RPE1 copy number classification and theoretical sensitivity after adding each probe until all 120 probes were incorporated. This progressive addition strategy allowed us to observe how sensitivity evolves as panel size increases, revealing both the rate of improvement and potential saturation points.

Mean theoretical sensitivity increased monotonically with probe number across all copy number levels, though the rate of improvement and asymptotic behavior varied by copy number state (**Fig S5C**). For the 5-component classification model (distinguishing 1, 2, 3, 4, and 5 copies), the simulation demonstrated that achieving at least 90% mean sensitivity requires 4 probes for 1-copy states, 20 probes for 2-copy states, 50 probes for 3-copy states, and 90 probes for 4-copy states. The 5-copy state proved most challenging, reaching a maximum mean sensitivity of approximately 87% even with the full 120-probe panel. The variability in sensitivity estimates across the 50 trials decreased progressively as probe number increased, indicating that larger panels provide not only higher accuracy but also more consistent performance.

For applications where precise copy number enumeration is less critical than distinguishing between loss, neutral, and gain states, we evaluated a simplified 3-component Gaussian mixture model representing 1 copy (loss), 2 copies (neutral), and ≥3 copies (gain). Under this classification scheme, sensitivity improvements were more pronounced (**Fig S5D**). The 90% sensitivity threshold was achieved at 4 probes for loss states, 20 probes for neutral states, and 26 probes for gain states. Furthermore, 99% sensitivity—representing near-perfect classification—could be attained with 22 probes for loss detection, 66 probes for neutral state classification, and 76 probes for gain detection. These results demonstrate that the choice of classification granularity substantially impacts probe requirements, with simpler classification schemes achieving high accuracy with fewer probes.

### Sequencing Depth Shows Minimal Impact on Classification Sensitivity

While probe number emerged as a critical determinant of classification performance, we also investigated whether sequencing depth—the average number of reads per cell per probe—significantly influences copy number calling accuracy. To evaluate this parameter, we systematically downsampled the read count matrix from our experimental dataset to 80%, 60%, 40%, and 20% of the original average depth (corresponding to approximately 68, 51, 34, and 17 reads per cell per probe, respectively, from the full depth of 85 reads). For each downsampling level, we generated 50 independent downsampled datasets and recalculated copy number classifications for RPE1 cells.

Across all copy number levels, the effect of reduced sequencing depth on theoretical sensitivity was modest: mean theoretical sensitivity decreased by an average of only 0.4 percentage points at 80% depth, 0.8 percentage points at 60% depth, 1.8 percentage points at 40% depth, and 4.4 percentage points at 20% depth relative to full sequencing depth. These marginal decreases indicate that, beyond a moderate threshold of approximately 35-50 reads per cell per probe, additional sequencing depth yields diminishing returns in terms of classification accuracy.

To formally test whether the effects of sequencing depth and probe number on sensitivity were independent or synergistic, we performed analysis of variance (ANOVA) on the combined simulation data. The lack of a significant interaction term (p = 0.9-1.0 for all copy number levels) confirmed that these two technical parameters influence classification performance through independent mechanisms. This independence has important practical implications: it suggests that optimizing probe number and sequencing depth can be approached as separate design considerations, with probe number being the primary lever for improving sensitivity and sequencing depth primarily affecting data quality and reliability rather than fundamental classification capability.

The minimal sensitivity gains beyond moderate sequencing depths also have favorable economic implications. Since sequencing costs scale directly with depth, these results indicate that cost-efficient experimental designs can achieve near-optimal classification performance without requiring excessive sequencing coverage. For most applications, targeting 40-60 reads per cell per probe should provide an optimal balance between classification accuracy and sequencing cost.

### Summary and Design Recommendations

Our systematic simulation analyses establish probe number as the dominant determinant of copy number classification sensitivity in KaryoTap, with sequencing depth playing a secondary role beyond moderate coverage thresholds. The relationship between probe number and sensitivity follows a logarithmic pattern, with diminishing returns as probe density increases but meaningful improvements achievable through strategic probe addition, particularly for smaller chromosomes and chromosome arms that are underrepresented in genome-proportional designs.

The choice between 5-component and 3-component classification models substantially impacts probe requirements, with the simpler 3-component model achieving high sensitivity with considerably fewer probes. For applications focused on detecting aneuploidy (presence or absence of gains/losses) rather than precise copy number enumeration, the 3-component model represents an efficient approach that maintains high sensitivity while minimizing sequencing costs.

These findings provide quantitative guidelines for designing future KaryoTap panels tailored to specific experimental objectives, balancing classification sensitivity against practical constraints of panel size and sequencing budget. The independence of probe number and sequencing depth effects further allows these parameters to be optimized separately, facilitating flexible experimental designs appropriate for diverse research applications.

### GMM-based Theoretical Accuracy (KaryoTap version V1)

The empirical accuracy for copy number calls in RPE1 only demonstrates the ability to detect 2 copies of a chromosome. We determined the theoretical accuracy for calling copy numbers of 1, 2, 3, 4, and 5 by calculating the proportion of each GMM component distribution that would be correctly classified. Overall, the highest theoretical accuracy was detected for 1 copy with an average of 97%, decreasing with each additional copy; 2 copies had an average sensitivity of 83% and 3 copies had 64% (**Fig S3E**). The theoretical sensitivity for 2 copies strongly correlated with the empirical accuracy for RPE1 calls (Pearson r = 0.97). As expected, theoretical sensitivity at all 5 copy number levels decreased for chromosomes with fewer probes (**Fig S3E**). Theoretical arm-level sensitivity averaged 95% for 1 copy, 74% for 2 copies, and 53% for 3 copies.

### Cost Comparison of KaryoTap compared to Similar Methods

To evaluate Karyotap compared to other single-cell genotyping methods, we compared KaryoTap to Arc-well data downsampled to 30K and 15K reads per cell. Downsampling to 15K reads showed qualitative increases in noisiness, with only 30-50% of full-depth events detected. In contrast, KaryoTap RPE1 data at comparable depth (∼15K reads per cell) maintained robust detection. In terms of cost, GenomePlex WGA4 (DOP-PCR WGA (60) costs $19.45–27.70 per cell, pricing a 500-cell experiment at $9,725 before sequencing. Alternative shallow scWGS methods based on cell capture in single wells cost approximately

$8-10 per cell (61). KaryoTap experiments cost approximately $0.5-1 per cell—less than 10% of these alternatives. By achieving comparable detection sensitivity at a fraction of the cost compared to WGS methods, KaryoTap provides a scalable and cost-effective solution for high-throughput aneuploidy detection that overcomes the size-dependent biases inherent to WGS-based approaches.

## List of Abbreviations

|  |  |  |
| --- | --- | --- |
| 1369 | BFP | Blue florescent protein |
| 1370 | CIN | Chromosomal instability |
| 1371 | CNV | Copy number variant |
| 1372 | DLP | Direct library preparation |
| 1373 | DOP-PCR | Degenerate-oligonucleotide-primed PCR |
| 1374 | FACS | Fluorescence associated cell sorting |
| 1375 | GMM | Gaussian mixture model |
| 1376 | gRNA | guide RNA |
| 1377 | hCECs | Human colorectal epithelial cells |
| 1378 | hPNEs | Human pancreatic nestin-expressing cells |
| 1379 | hMEC | Human mammary epithelial cells |
| 1380 | HMEL | Human Melanocytes |
| 1381 | hPDE | Human pancreatic duct epithelial cells |
| 1382 | MALBAC | Multiple annealing and looping-based amplification cycles |
| 1383 | MDA | Multiple displacement amplification |
| 1384 | MOI | Multiplicity of infection |
| 1385 | ORF | Open reading frame |
| 1386 | PCA | Principal components analysis |
| 1387 | PCR | Polymerase chain reaction |
| 1388 | PDF | Probability Density Function |
| 1389 | RPE1 | Retinal pigment epithelial (cells) 1 |
| 1390 | scDNA-seq | Single-cell DNA sequencing |
| 1391 | scRNA-seq | Single-cell RNA sequencing |
| 1392 | sgRNA | Single guide RNA |
| 1393 | SNP | Single-nucleotide polymorphism |
| 1394 | SNV | Single nucleotide variant |
| 1395 | VAF | Variant allele frequency |
| 1396 | WGA | Whole genome amplification |
| 1397 | WGS | Whole genome sequencing |
| 1398 |  |  |
| 1399 |  |  |

## Declarations

### Ethics Approval and Consent to Participate

Not applicable

### Consent for Publication

Not applicable

### Availability of Data and Materials

Sequencing data for Tapestri experiments are available in the SRA repository under NCBI BioProject accession PRJNA950110, https://www.ncbi.nlm.nih.gov/bioproject/950110 (53). The karyotapR package is available on GitHub at http://github.com/joeymays/karyotapR (54).The source code for karyptapR version 0.1 used for this study is archived on Zenodo under DOI https://doi.org/10.5281/zenodo.8305561(55). All data analysis scripts used in this study are available at https://github.com/joeymays/karyotap-publication (56) and are archived on Zenodo under DOIhttps://doi.org/10.5281/zenodo.8329277. Tapestri Pipeline output files used in this study are available on Zenodo under DOI https://doi.org/10.5281/zenodo.8305841(57).

### Authors’ contributions

JCM designed the study, performed wet lab experiments and bioinformatics analyses, developed the karyotapR software package, and wrote the manuscript for the KaryoTap development. FA design and performed experiments and bioinformatics analyses and wrote the manuscript for the in vitro screens. MT helped with the LOH analysis, experimental design and edited the manuscript. SM performed lentiviral preparation and transductions. NB and XZ designed Tapestri panels. JJB and HQ performed and supported cell culture. GRK prepared barcode vectors. LJH supervised research. TD designed the study, edited the manuscript, and supervised research.

## Acknowledgments

NYU Langone’s Genome Technology Center (RRID: SCR_017929) is supported by the Cancer Center Support Grant P30CA016087 at the Laura and Isaac Perlmutter Cancer Center. The computational requirements for this work were supported in part by the NYU Langone High Performance Computing (HPC) Core’s resources and personnel. We thank Paolo Mita, Pan Cheng, and Lisa Katsnelson for their helpful suggestions and insights in the preparation of this work.

## Additional Files

● AdditionalFile01.csv: Probe Design for Panel Version 1 (CO216)
● AdditionalFile02.csv: Probe Design for Panel Version 2 (CO610)
● AdditionalFile03.csv: Probe Design for Panel Version 3 (CO810)

## Supplementary Table Legends

Supplementary Table 1

Number of probes per chromosome arm for custom Tapestri panels.

Supplementary Table 2

Theoretical sensitivity of panel Version 1 copy number calls for each chromosome/chromosome arm at copy number values of 1, 2, 3, 4 and 5, calculated from GMMs.

Supplementary Table 3

Theoretical sensitivity of panel Version 2 copy number calls for each chromosome/chromosome arm at copy number values of 1, 2, 3, 4 and 5, calculated from 5-component GMMs. “Gain” column indicates theoretical sensitivity for detecting chromosomal gains or 23 copies using 3-component GMM. 1 copy (5-component model) and “loss” detection (3-component model) sensitivities are equivalent. 2 copy (5-component model) and “neutral” detection (3-component model) sensitivities are equivalent.

Supplementary Table 4

Sequences of barcodes and gRNAs.

Supplementary Table 5

Results of Fisher’s exact test for chromosomal gains and losses induced by KaryoCreate. p-values from Fisher’s Exact test comparing the proportion of copy number gains and losses in each chromosome in each sample to the corresponding chromosome in the sgNC negative control sample are given.

Supplementary Table 6

Percentage of chromosomal gains and losses Induced by KaryoCreate. Percentage indicates the proportion of cells in each KayroCreate-treated sample with either 1, 2, or 3+ copies of each chromosome.

Supplementary Table 7

Theoretical sensitivity of panel Version 3 copy number calls for each chromosome/chromosome arm at copy number values of 1, 2, 3, 4 and 5, calculated from 5-component GMMs. “Gain” column indicates theoretical sensitivity for detecting chromosomal gains or 23 copies using 3-component GMM. 1 copy (5-component model) and “loss” detection (3-component model) sensitivities are equivalent. 2 copy (5-component model) and “neutral” detection (3-component model) sensitivities are equivalent.

Supplementary Table 8

Sequences of barcodes used in the aneuploidy induction and in vitro screening experiment.

---

**Supplementary Figure 1.**
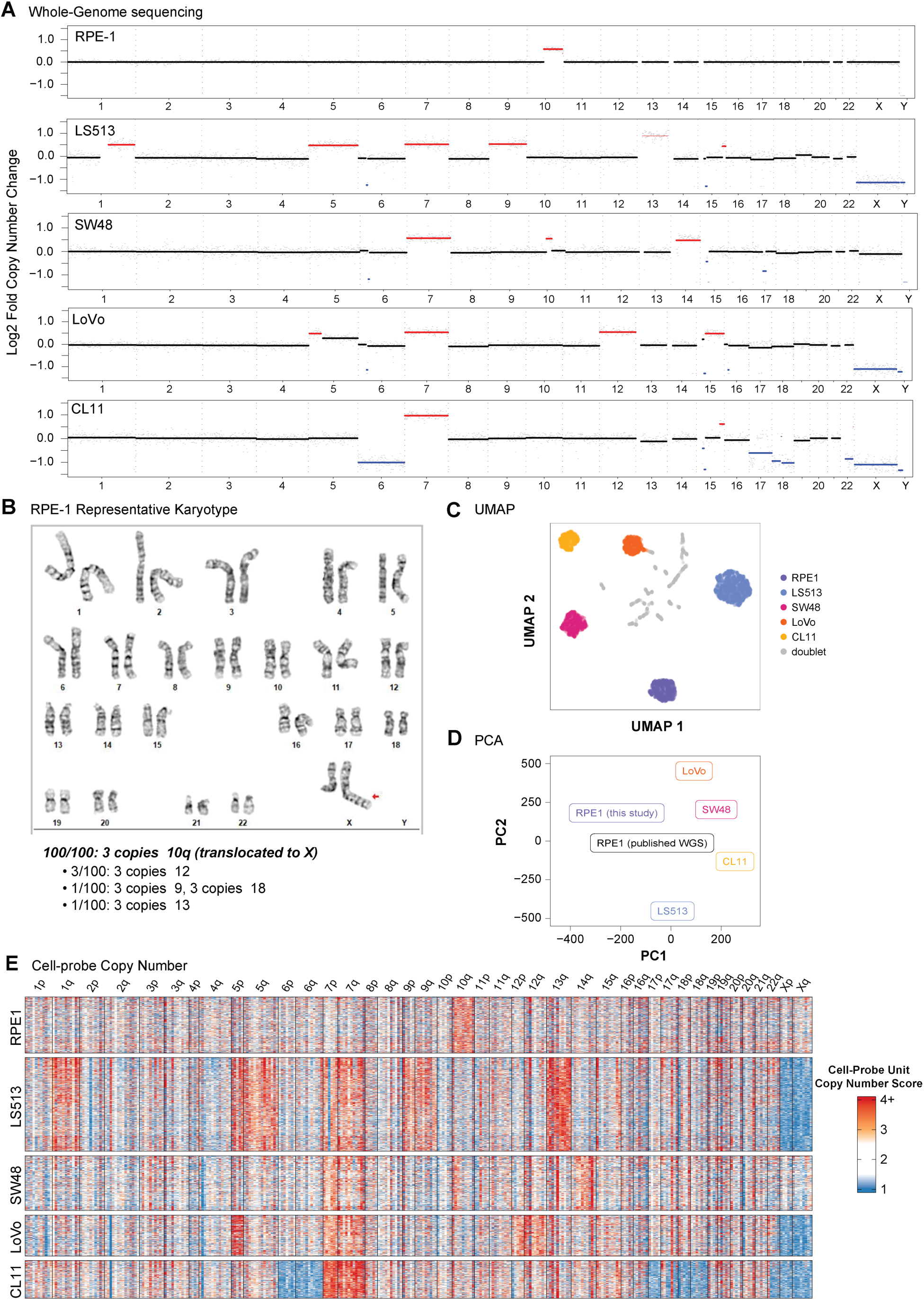
(A) Bulk low-pass whole genome sequencing of RPE1, LS513, SW48, LoVo, and CL11 cell lines. Red highlight indicates amplification of at least one copy of highlighted segment; blue similarly indicates deletion. **(B)** Representative g-banded karyogram of RPE1, indicating additional copy of chr10q translocated to the X chromosome (red arrow). **(C)** UMAP projection of top 4 principal components of allele frequencies for N=2,986 cells representing 5 cell lines. Clustering was performed using the dbscan method. Cells were considered doublets if they were not members of the 5 largest clusters. **(D)** PCA plot of first two principal components of mean allele frequencies for previously published deep sequencing of RPE1 and the 5 cell lines analyzed by scDNA-seq. **(E)** Heatmap of cell-probe copy number values for five cell lines using custom Tapestri panel Version 1. Probes are organized by chromosome arm in genomic order.

**Supplementary Figure 2.**
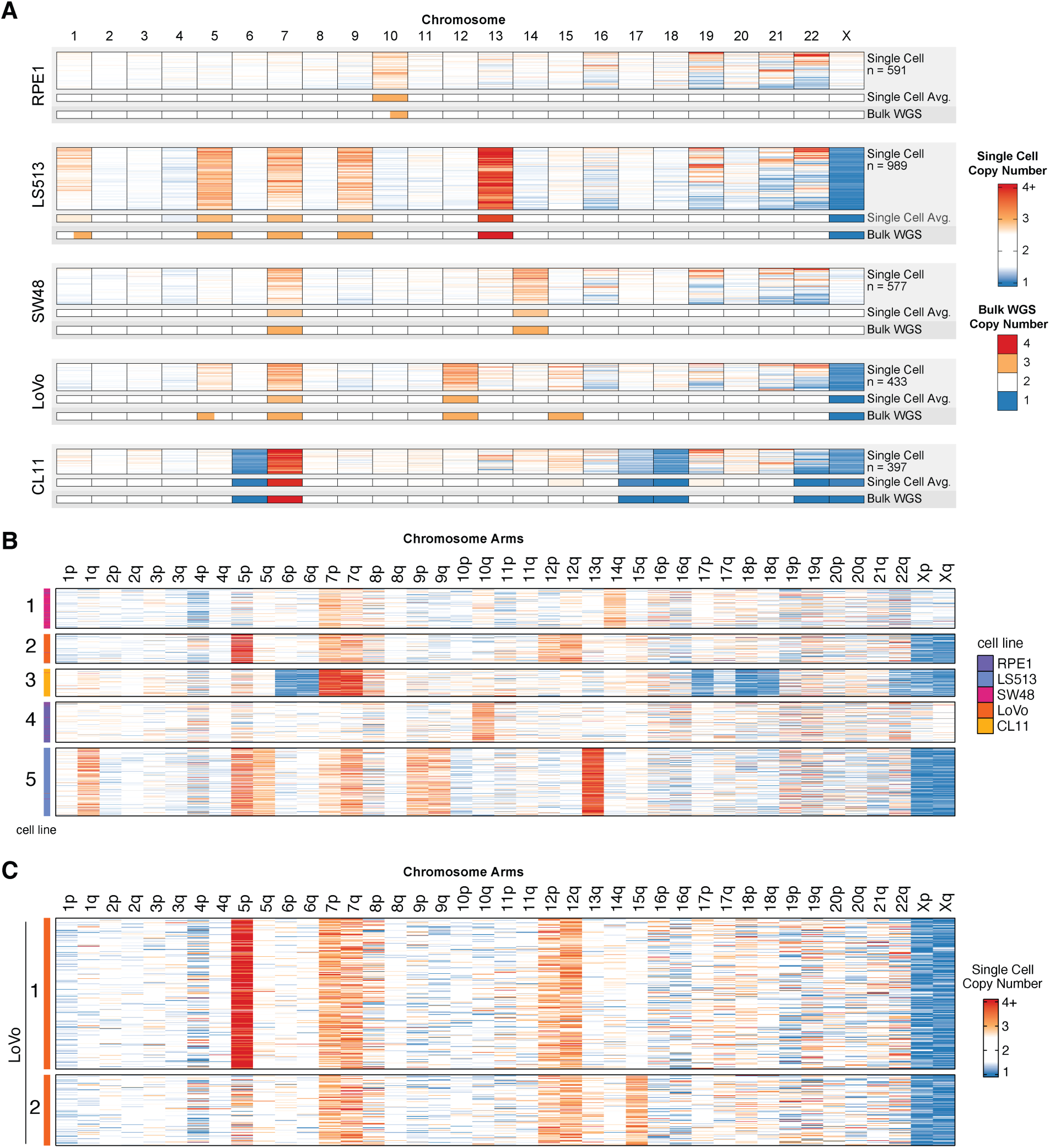
**(A)** Heatmap of copy number scores for each cell-chromosome unit for 5 cell lines using custom Tapestri panel Version 1. Upper blocks indicate copy number scores for each cell, middle blocks indicate average intensity of upper blocks, and lower blocks indicate copy number from bulk WGS (see Fig S1A). Half-filled lower blocks indicate chromosome arm-level aneuploidy. Number of cells included in single-cell blocks is indicated. **(B)** Heatmap of GMM calls for chromosome arms across 5 cell lines using v1 panel. **(C)** Heatmap for further clustering within LoVo.

**Supplementary Figure 3.**
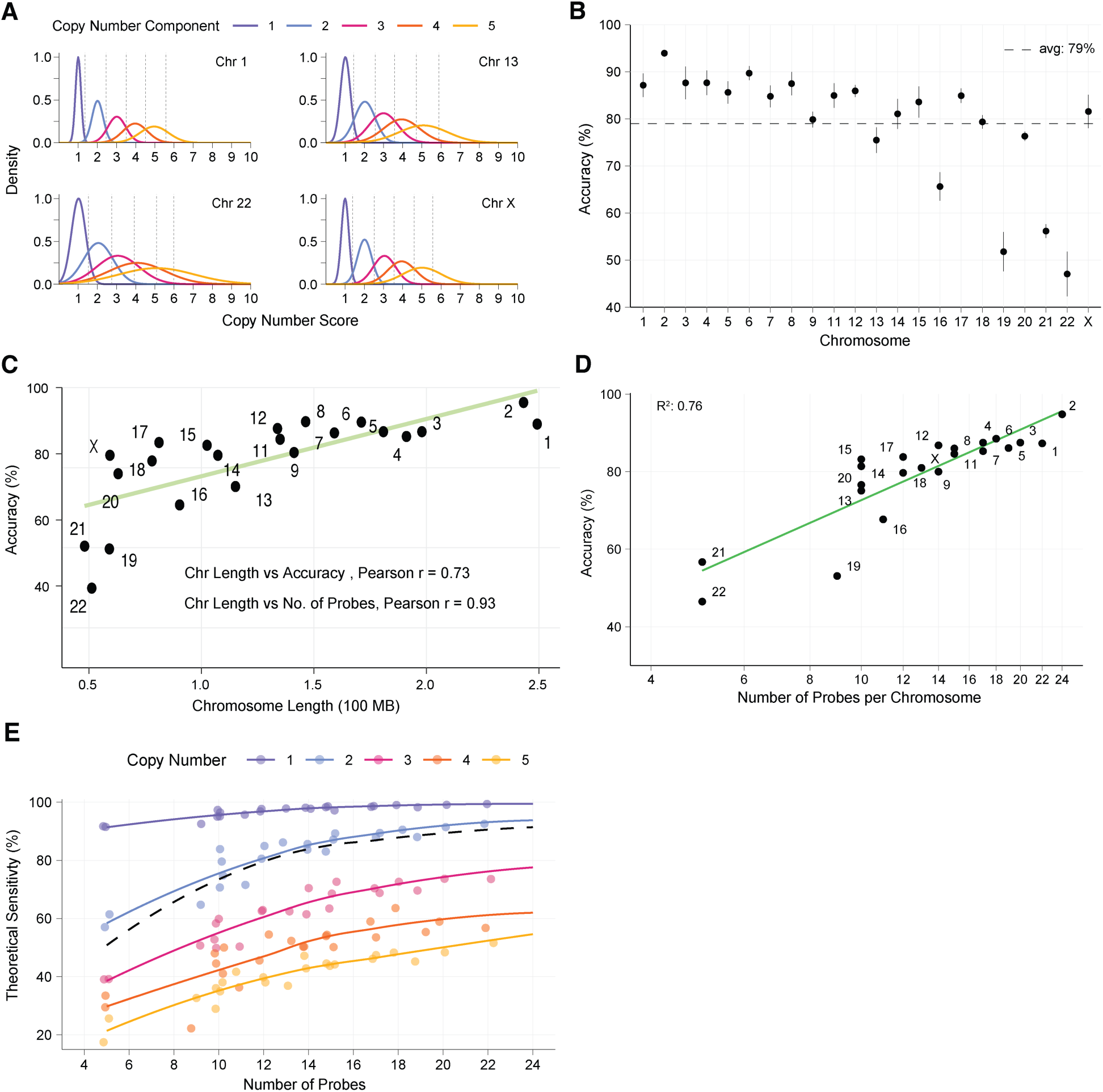
**(A)** Probability density functions of Gaussian mixture models (GMM) fit for chromosomes 1, 13, 22, and X using RPE1 cells. Dotted lines indicate decision boundaries between GMM components. **(B)** 5-fold cross validation of copy number call accuracy for RPE1 cells by chromosome. Chr10 is omitted. Dot indicates mean accuracy, and lines indicate ± mean absolute deviation. Horizontal dotted line indicates average (avg) accuracy across chromosomes. **(C)** Chromosome length (in 100 megabases) vs. accuracy of RPE1 panel Version 1 copy number calls chromosome. Chr10 is omitted. Trendline fit by linear regression. **(D)** Correlation between number of probes per chromosome and karyotap chromosome-copy-number call accuracy. **(E)** Theoretical copy number call sensitivity for each chromosome and copy number level calculated from GMMs. Points are slightly jittered horizontally to decrease overlapping.

**Supplementary Figure 4.**
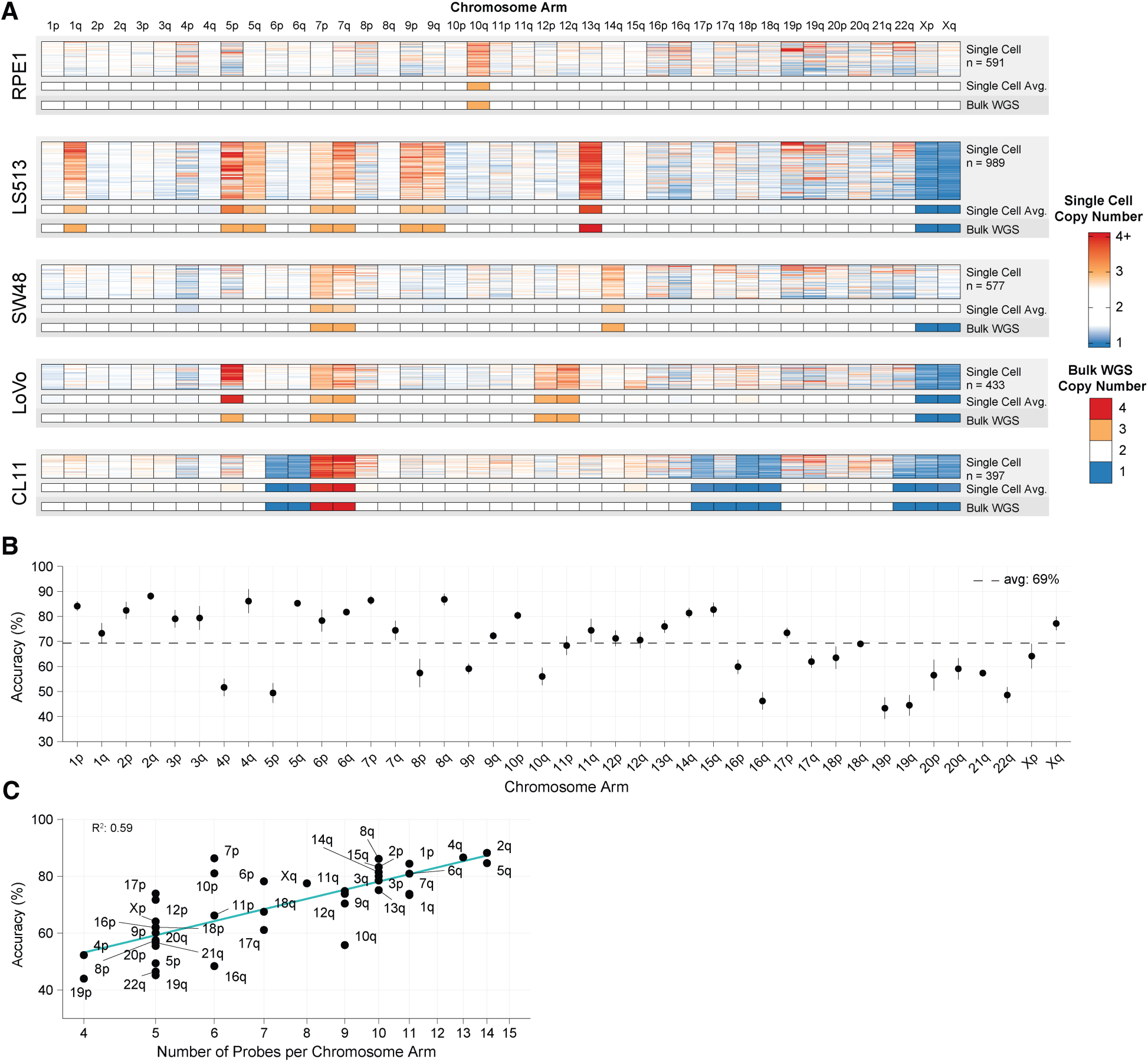
**(A)** Heatmap of copy number scores for each cell, smoothed across chromosome arms, for five cell lines using custom Tapestri panel Version 1. Upper blocks indicate copy number scores for each cell; middle blocks indicate average intensity of upper blocks, and lower blocks indicate copy number from bulk WGS (see Fig S1A). **(B)** 5-fold cross validation of copy number call accuracy for RPE1 cells by chromosome arm. Dot indicates mean accuracy and lines indicate ± mean absolute deviation. Horizontal dotted line indicates average accuracy across chromosome arms. **(C)** Linear regression of RPE1 copy number call accuracy for each chromosome arm on number of probes per chromosome arm. X-axis is log-scaled to reflect log transformation of number of probes in regression.

**Supplementary Figure 5.**
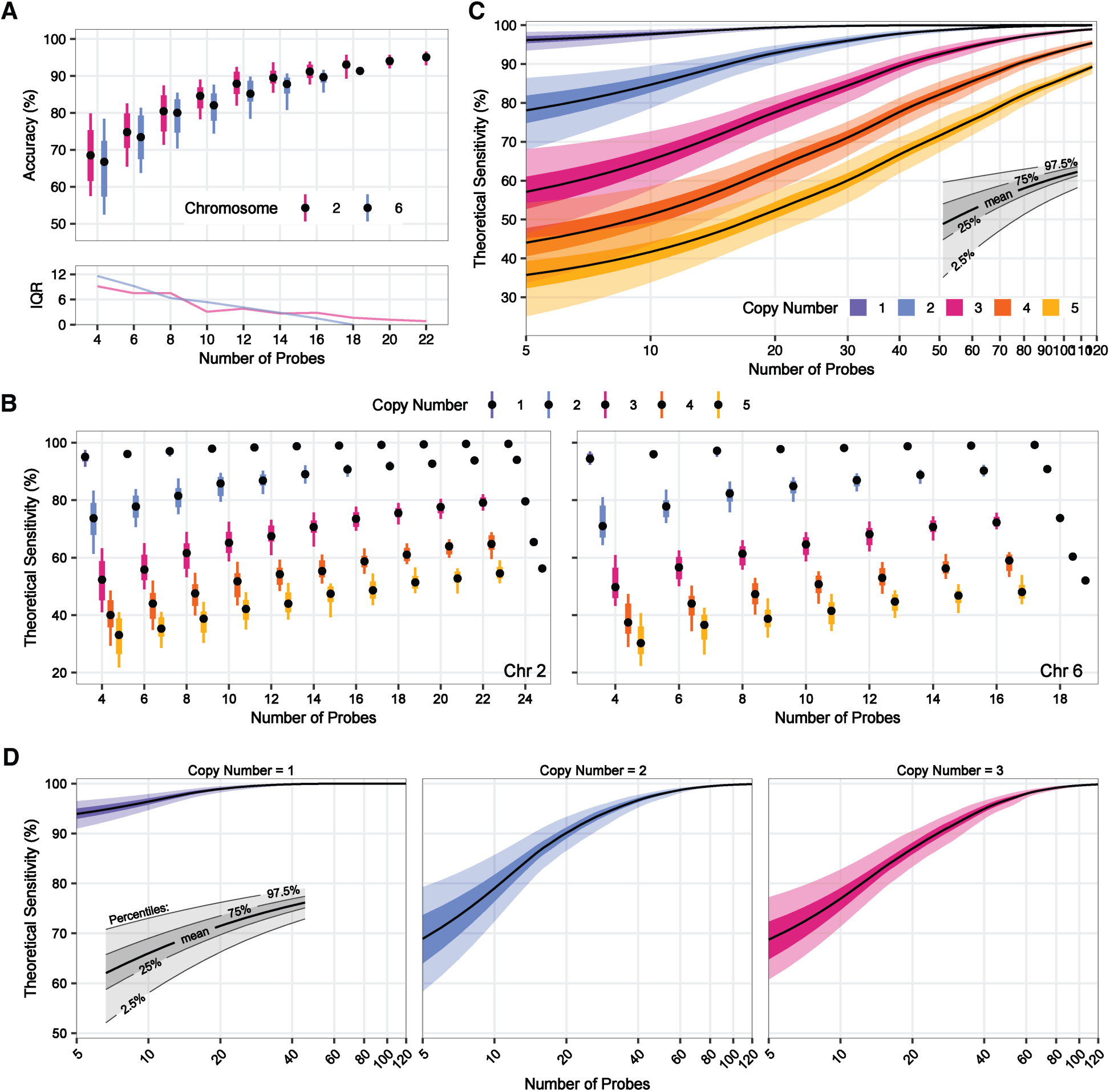
Effects on copy number call accuracy on probe sampling simulations. **(A)** Box plots and interquartile range (IQR) of accuracy from 50 probe downsampling simulations for chr2 and chr6.Boxes encompass middle 50%, whiskers encompass middle 95%, dot indicates median. **(B)** Theoretical sensitivity of 50 panel simulations. Values for each copy number level were smoothed by Loess regression. The black line represents the mean, the darker inner shading indicates the middle 50% of the data, and the lighter outer shading represents the middle 95% of the data. **(C)** Theoretical sensitivity of 50 panel simulations. Values for each copy number level were smoothed by Loess regression. The black line represents the mean, the darker inner shading indicates the middle 50% of the data, and the lighter outer shading represents the middle 95% of the data. **(D)** Theoretical copy number call sensitivity of 50 panel simulations, using a 3 component GMM. Values for each copy number level were smoothed by Loess regression. The black line represents the mean, the darker inner shading indicates the middle 50% of the data, and the lighter outer shading represents the middle 95% of the data.

**Supplementary Figure 6.**
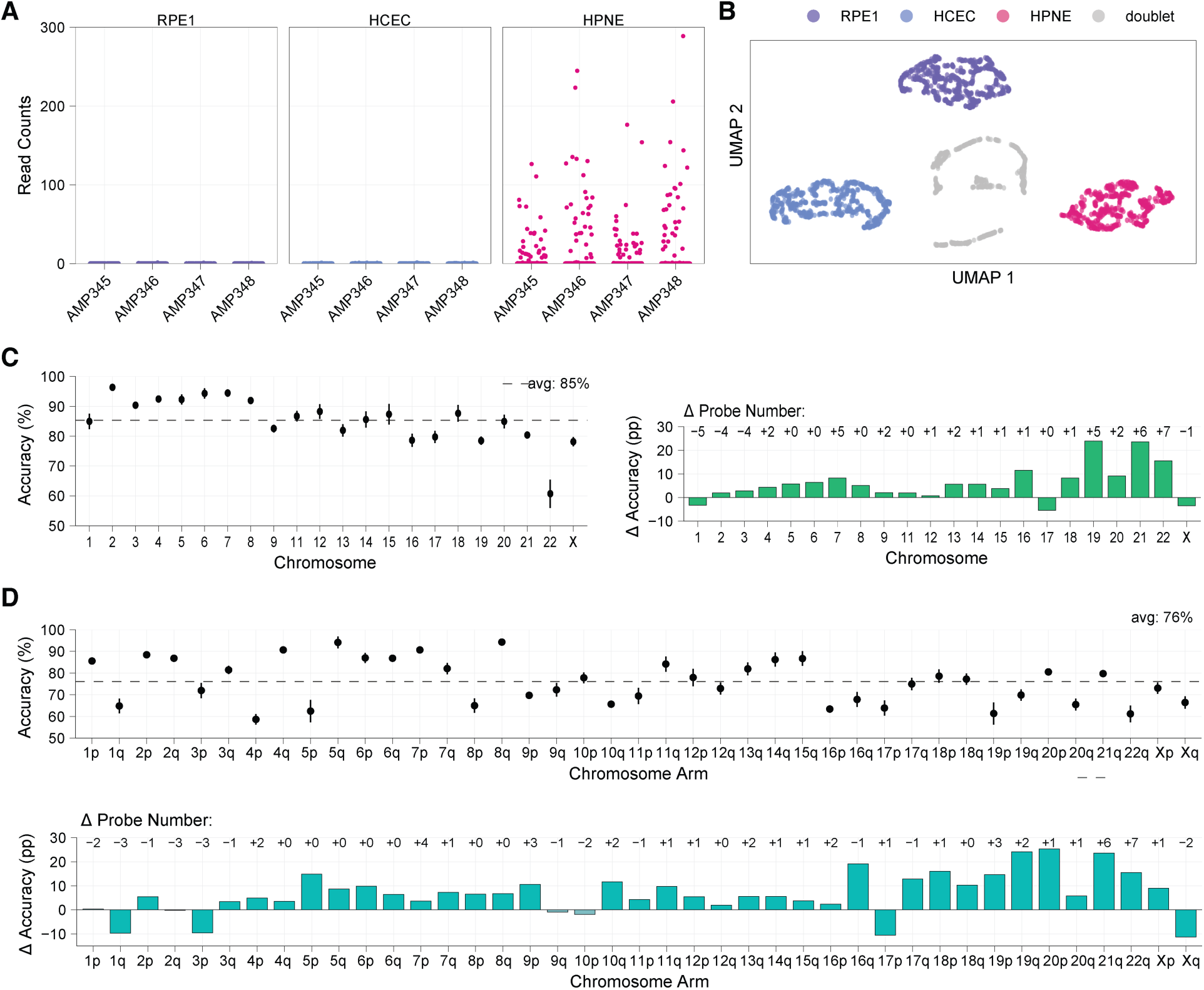
**(A)** Read counts per cell of four probes targeting chrY across 3 cell lines. **(B)** UMAP projection of top 2 principal components of allele frequencies for N=2,347 cells representing 3 cell lines. Clustering was performed using the dbscan method. Cells were considered doublets if they were not members of the 3 largest clusters. **(C)** 5-fold cross validation of copy number call sensitivity for RPE1 cells by chromosome using panel Version 2. Chr10 is omitted. Dot indicates mean accuracy, and lines indicate ± mean absolute deviation. Horizontal dotted line indicates average (avg) accuracy across chromosomes. On the right, the barplot shows the change in copy number call accuracy by chromosome for RPE1 cells between Version 2 and Version 1 panels. Δ Probe Number is the difference in number of probes targeting a given chromosome between Version 2 and Version 1. pp: percentage points. **(D)** 5-fold cross validation of copy number call accuracy for RPE1 cells by chromosome arms using custom Tapestri panel Version 2. Dot indicates mean accuracy, and lines indicate ± mean absolute deviation. Horizontal dotted line indicates average accuracy across chromosome arms. On the right, the barplot shows the change in copy number call accuracy by chromosome arm for RPE1 cells between Version 2 and Version 1 panels. Δ Probe Number is the difference in number of probes targeting a given chromosome arms between Version 2 and Version 1. pp: percentage points.

**Supplementary Figure 7.**
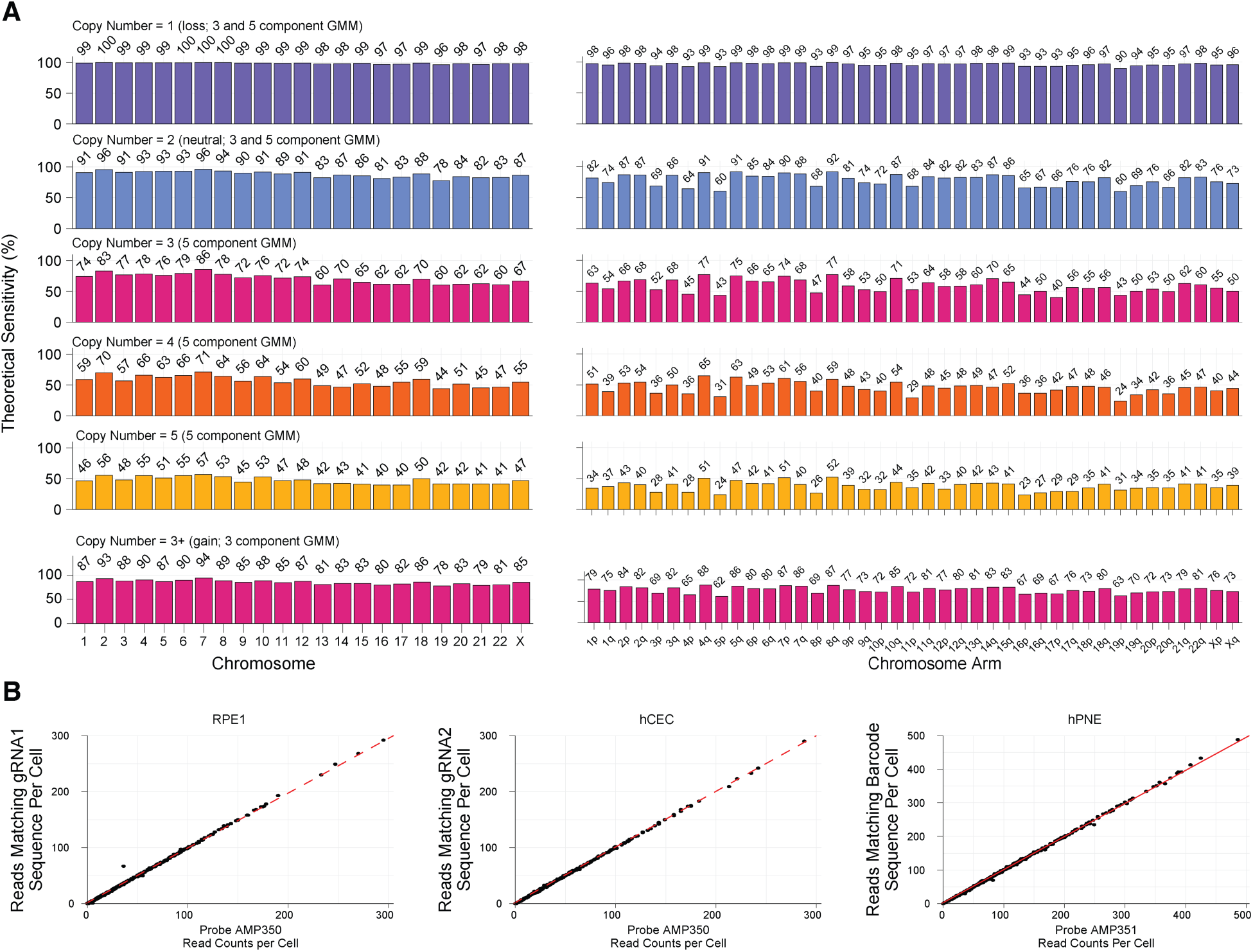
(B) Mean theoretical accuracy of panel Version 3 copy number calls for each chromosome at copy number values of 1, 2, 3, 4 and 5. **(C)** Relationship between read counts per cell for either the gRNA or DNA Barcode probes (x-axis) and the number of reads per cell that match the specific gRNA or DNA Barcode sequences (y-axis). Dotted red line indicates *x* = *y*.

**Supplementary Figure 8.**
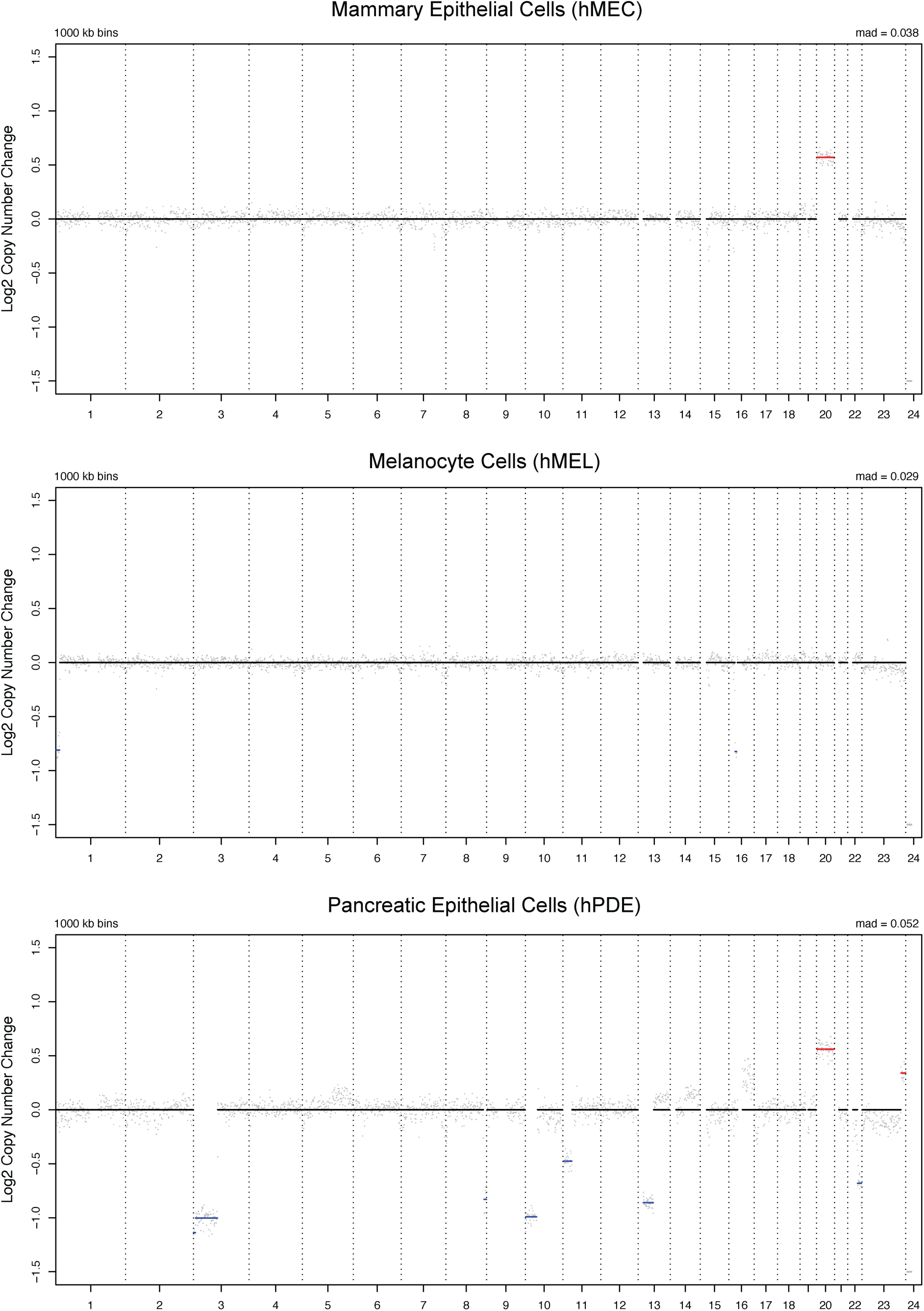
Low-pass WGS profile for mammary epithelial cells (hMEC), melanocyte cells (hMEL) and pancreatic epithelial cells (hPDE).

**Supplementary Figure 9.**
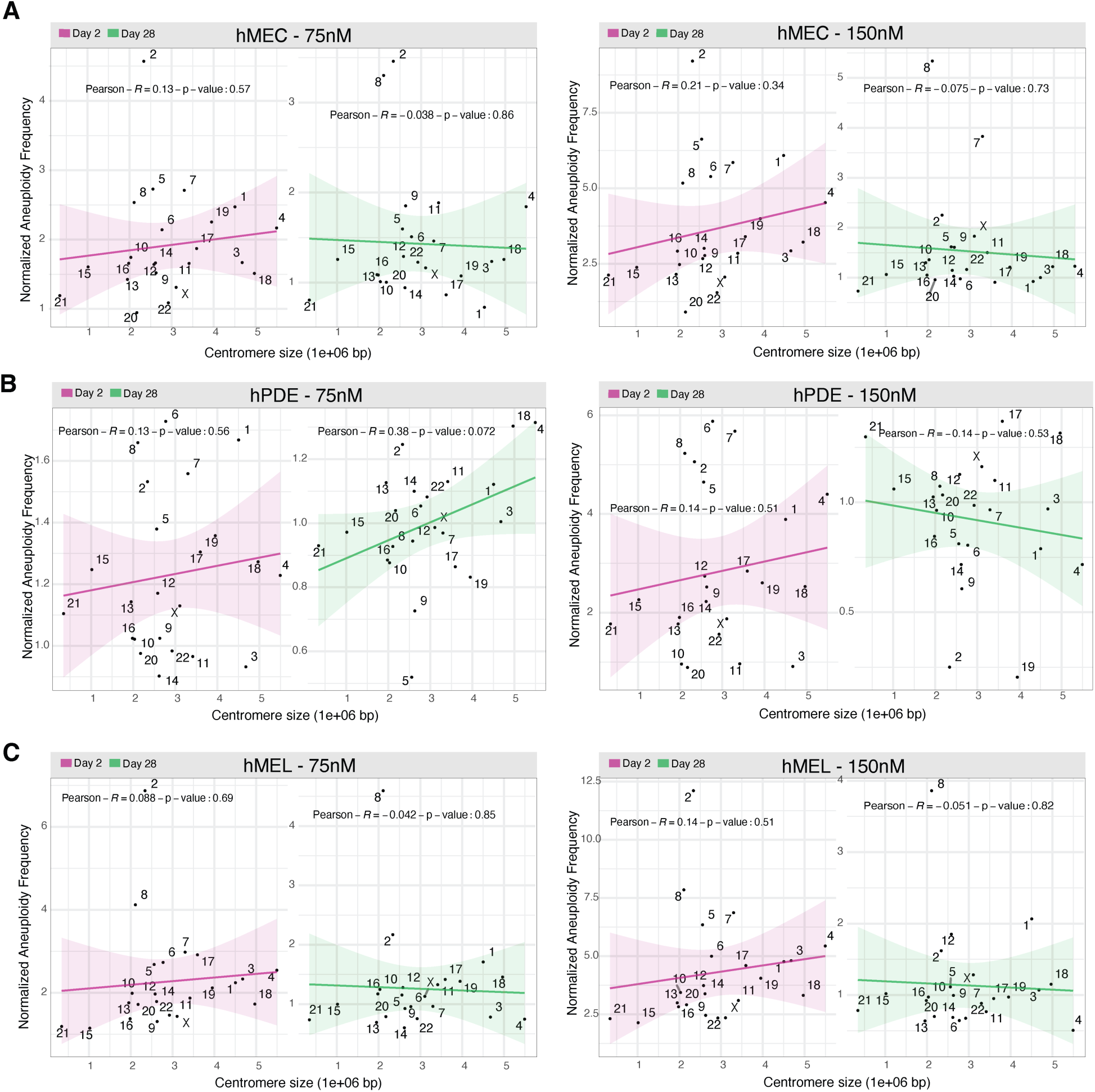
Correlation between centromere size and aneuploidy frequency (gains and losses) following reversine treatment for **(A)** mammary epithelial cells (hMEC), **(B)** melanocyte cells (hMEL) and **(C)** pancreatic epithelial cells (hPDE). Regression lines represent trends for 75 nM and 150 nM reversine treatments; correlation coefficients (Pearson’s R) and p-values are provided for statistical significance.

**Supplementary Figure 10.**
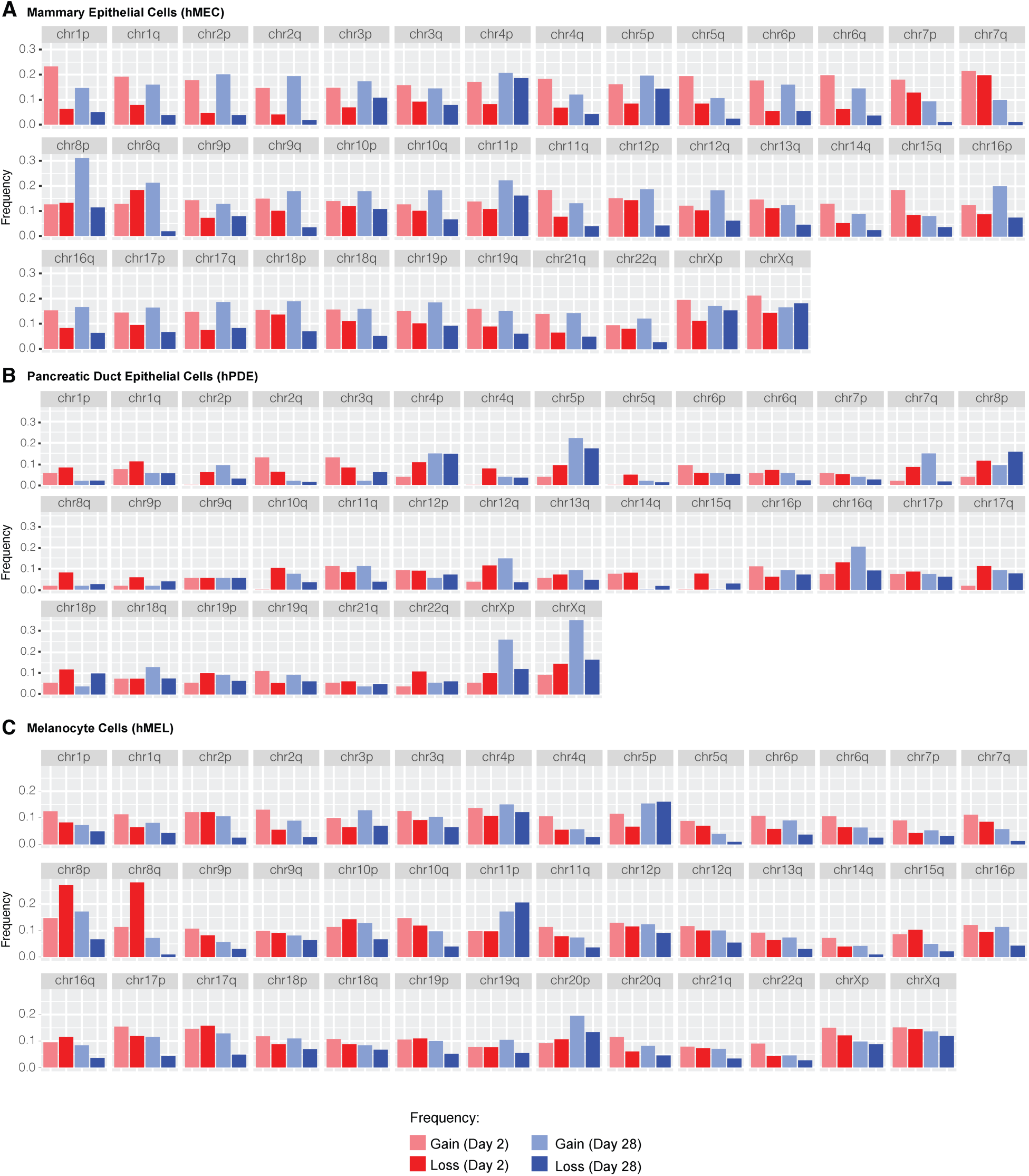
Frequency of gains and losses across all chromosomal arms at the two time points, Day 2 and Day 28 for hMEC **(A)**, hPDE **(B)**, and hMEL **(C)**.

**Supplementary Figure 11.**
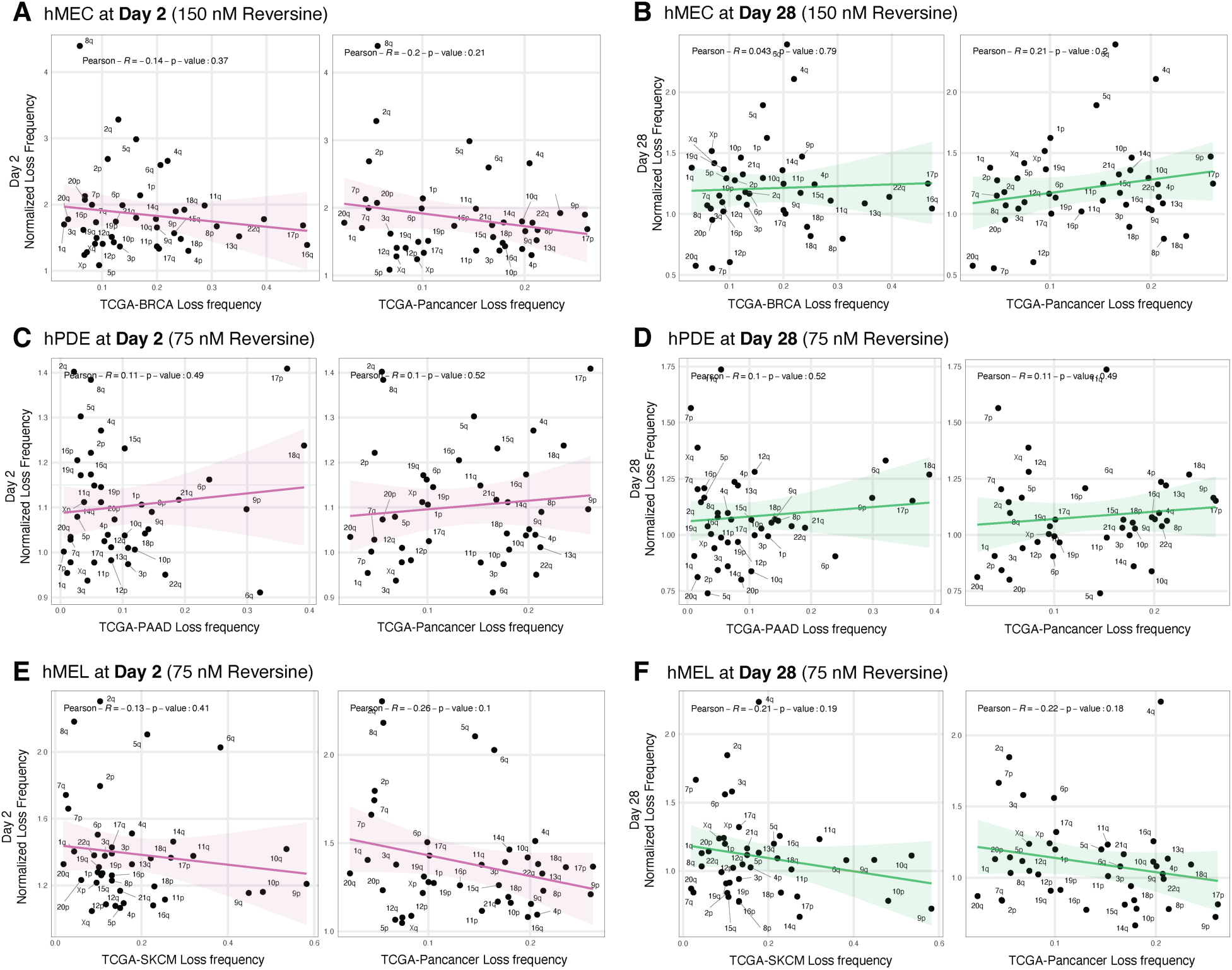
Comparison of chromosome-arm loss frequencies induced by reversine with cancer amplification frequency from TCGA data. Panels compare normalized chromosome arm loss frequencies at Day 2 (A, C, E) and Day 28 (B, D, F) with TCGA deletion frequency from respective cancer types (hMEC with BRCA, hPDE with PAAD, and hMEL with SKCM) and Pancancer. Scatter plots include correlation coefficients (Pearson’s R) and p-values. Green and pink regression lines distinguish between gains at Day 2 (pink) and Day 28 (green).

**Supplementary Figure 12.**
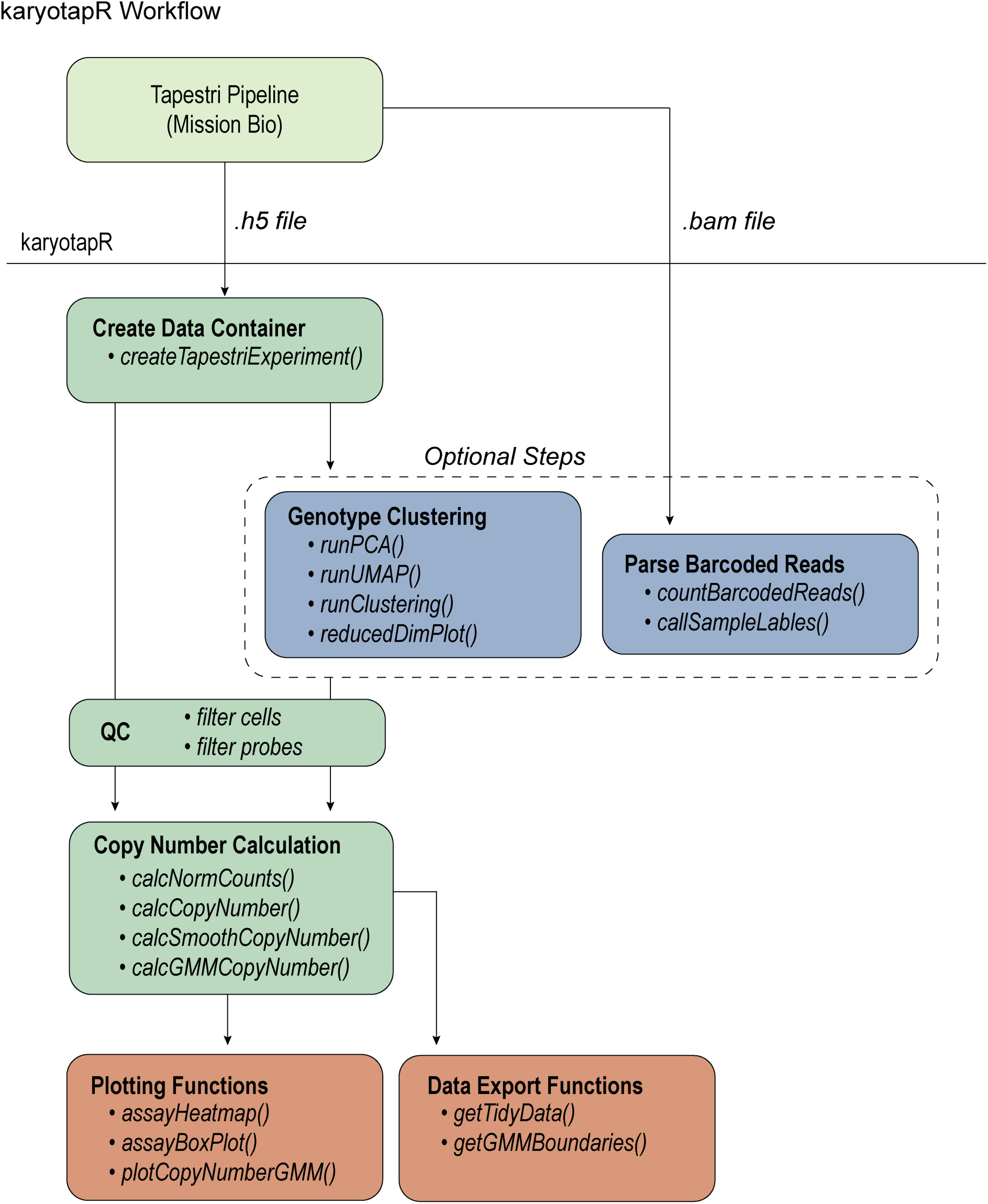
Overview of workflow for copy number analysis using karyotapR R package.

## References

1. Navin N, Krasnitz A, Rodgers L, Cook K, Meth J, Kendall J, et al. Inferring tumor progression from genomic heterogeneity. Genome Res. 2010 Jan;20(1):68–80.

2. Ben-David U, Amon A. Context is everything: aneuploidy in cancer. Nature Reviews Genetics. 2020 Jan;21(1):44–62.

3. Weaver BAA, Cleveland DW. Aneuploidy: Instigator and Inhibitor of Tumorigenesis. Cancer Research. 2007 Nov 1;67(21):10103–5.

4. Bakhoum SF, Ngo B, Laughney AM, Cavallo JA, Murphy CJ, Ly P, et al. Chromosomal instability drives metastasis through a cytosolic DNA response. Nature. 2018 Jan;553(7689):467–72.

5. Bakhoum SF, Cantley LC. The Multifaceted Role of Chromosomal Instability in Cancer and Its Microenvironment. Cell. 2018 Sep 6;174(6):1347–60.

6. Ippolito MR, Martis V, Martin S, Tijhuis AE, Hong C, Wardenaar R, et al. Gene copy-number changes and chromosomal instability induced by aneuploidy confer resistance to chemotherapy. Developmental Cell. 2021 Sep 13;56(17):2440–2454.e6.

7. Lee AJX, Endesfelder D, Rowan AJ, Walther A, Birkbak NJ, Futreal PA, et al. Chromosomal Instability Confers Intrinsic Multidrug Resistance. Cancer Research. 2011 Mar 1;71(5):1858–70.

8. Anagnostou V, Smith KN, Forde PM, Niknafs N, Bhattacharya R, White J, et al. Evolution of Neoantigen Landscape during Immune Checkpoint Blockade in Non–Small Cell Lung Cancer. Cancer Discovery. 2017 Mar 5;7(3):264–76.

9. Lynch AR, Bradford S, Zhou AS, Oxendine K, Henderson L, Horner VL, et al. A survey of CIN measures across mechanistic models [Internet]. bioRxiv; 2023 [cited 2023 Aug 16]. p. 2023.06.15.544840. Available from: https://www.biorxiv.org/content/10.1101/2023.06.15.544840v1

10. Bakker B, Taudt A, Belderbos ME, Porubsky D, Spierings DCJ, de Jong TV, et al. Single-cell sequencing reveals karyotype heterogeneity in murine and human malignancies. Genome Biology. 2016 May 31;17(1):115.

11. Baslan T, Hicks J. Unravelling biology and shifting paradigms in cancer with single-cell sequencing. Nat Rev Cancer. 2017 Sep;17(9):557–69.

12. Navin N, Kendall J, Troge J, Andrews P, Rodgers L, McIndoo J, et al. Tumour evolution inferred by single-cell sequencing. Nature. 2011 Apr;472(7341):90–4.

13. Wang Y, Waters J, Leung ML, Unruh A, Roh W, Shi X, et al. Clonal evolution in breast cancer revealed by single nucleus genome sequencing. Nature. 2014 Aug;512(7513):155–60.

14. Bosco N, Goldberg A, Zhao X, Mays JC, Cheng P, Johnson AF, et al. KaryoCreate: A CRISPR-based technology to study chromosome-specific aneuploidy by targeting human centromeres. Cell [Internet]. 2023 Apr 18 [cited 2023 Apr 20]; Available from: https://www.sciencedirect.com/science/article/pii/S0092867423003264

15. Truong MA, Cané-Gasull P, de Vries SG, Nijenhuis W, Wardenaar R, Kapitein LC, et al. A kinesin-based approach for inducing chromosome-specific mis-segregation in human cells. The EMBO Journal. 2023 May 15;42(10):e111559.

16. Tovini L, Johnson SC, Guscott MA, Andersen AM, Spierings DCJ, Wardenaar R, et al. Targeted assembly of ectopic kinetochores to induce chromosome-specific segmental aneuploidies. The EMBO Journal. 2023 May 15;42(10):e111587.

17. Telenius H, Carter NP, Bebb CE, Nordenskjöld M, Ponder BAJ, Tunnacliffe A. Degenerate oligonucleotide-primed PCR: General amplification of target DNA by a single degenerate primer. Genomics. 1992 Jul 1;13(3):718–25.

18. Dean FB, Hosono S, Fang L, Wu X, Faruqi AF, Bray-Ward P, et al. Comprehensive human genome amplification using multiple displacement amplification. Proceedings of the National Academy of Sciences. 2002 Apr 16;99(8):5261–6.

19. Zong C, Lu S, Chapman AR, Xie XS. Genome-Wide Detection of Single-Nucleotide and Copy-Number Variations of a Single Human Cell. Science. 2012 Dec 21;338(6114):1622–6.

20. Mallory XF, Edrisi M, Navin N, Nakhleh L. Methods for copy number aberration detection from single-cell DNA-sequencing data. Genome Biology. 2020 Aug 17;21(1):208.

21. Zahn H, Steif A, Laks E, Eirew P, VanInsberghe M, Shah SP, et al. Scalable whole-genome single-cell library preparation without preamplification. Nat Methods. 2017 Feb;14(2):167–73.

22. Yin Y, Jiang Y, Lam KWG, Berletch JB, Disteche CM, Noble WS, et al. High-Throughput Single-Cell Sequencing with Linear Amplification. Molecular Cell [Internet]. 2019 Sep 5 [cited 2019 Sep 6]; Available from: http://www.sciencedirect.com/science/article/pii/S1097276519306185

23. Laks E, Zahn H, Lai D, McPherson A, Steif A, Brimhall J, et al. Resource: Scalable whole genome sequencing of 40,000 single cells identifies stochastic aneuploidies, genome replication states and clonal repertoires [Internet]. bioRxiv; 2018 [cited 2023 Jan 10]. p. 411058. Available from: https://www.biorxiv.org/content/10.1101/411058v2

24. Evrony GD, Hinch AG, Luo C. Applications of Single-Cell DNA Sequencing. Annu Rev Genomics Hum Genet. 2021 Aug 31;22:171–97.

25. Mission Bio. Mission Bio Tapestri. 2020 [cited 2020 May 7]. Tapestri: The Precision Genomics Platform. Available from: https://missionbio.com/tapestri/

26. Gawad C, Koh W, Quake SR. Single-cell genome sequencing: current state of the science. Nat Rev Genet. 2016 Mar;17(3):175–88.

27. Crasta K, Ganem NJ, Dagher R, Lantermann AB, Ivanova EV, Pan Y, et al. DNA breaks and chromosome pulverization from errors in mitosis. Nature. 2012 Feb;482(7383):53–8.

28. Reijns MAM, Parry DA, Williams TC, Nadeu F, Hindshaw RL, Rios Szwed DO, et al. Signatures of TOP1 transcription-associated mutagenesis in cancer and germline. Nature. 2022 Feb;602(7898):623–31.

29. Aird D, Ross MG, Chen WS, Danielsson M, Fennell T, Russ C, et al. Analyzing and minimizing PCR amplification bias in Illumina sequencing libraries. Genome Biology. 2011 Feb 21;12(2):R18.

30. Jothilakshmi S, Gudivada VN. Chapter 10 - Large Scale Data Enabled Evolution of Spoken Language Research and Applications. In: Gudivada VN, Raghavan VV, Govindaraju V, Rao CR, editors. Handbook of Statistics [Internet]. Elsevier; 2016 [cited 2023 Aug 22]. p. 301–40. (Cognitive Computing: Theory and Applications; vol. 35). Available from: https://www.sciencedirect.com/science/article/pii/S0169716116300463

31. Knudson AG. Mutation and Cancer: Statistical Study of Retinoblastoma. Proceedings of the National Academy of Sciences. 1971 Apr;68(4):820–3.

32. Shah JB, Pueschl D, Wubbenhorst B, Fan M, Pluta J, D’Andrea K, et al. Analysis of matched primary and recurrent BRCA1/2 mutation-associated tumors identifies recurrence-specific drivers. Nat Commun. 2022 Nov 7;13(1):6728.

33. Ryland GL, Doyle MA, Goode D, Boyle SE, Choong DYH, Rowley SM, et al. Loss of heterozygosity: what is it good for? BMC Medical Genomics. 2015 Aug 1;8(1):45.

34. Dong X, Zhang L, Hao X, Wang T, Vijg J. SCCNV: A Software Tool for Identifying Copy Number Variation From Single-Cell Whole-Genome Sequencing. Frontiers in Genetics [Internet]. 2020 [cited 2023 Jun 2];11. Available from: https://www.frontiersin.org/articles/10.3389/fgene.2020.505441

35. Shalem O, Sanjana NE, Zhang F. High-throughput functional genomics using CRISPR–Cas9. Nat Rev Genet. 2015 May;16(5):299–311.

36. Ly P, Eskiocak U, Kim SB, Roig AI, Hight SK, Lulla DR, et al. Characterization of Aneuploid Populations with Trisomy 7 and 20 Derived from Diploid Human Colonic Epithelial Cells. Neoplasia. 2011 Apr 1;13(4):348-IN17.

37. Sherry ST, Ward MH, Kholodov M, Baker J, Phan L, Smigielski EM, et al. dbSNP: the NCBI database of genetic variation. Nucleic Acids Res. 2001 Jan 1;29(1):308–11.

38. Nassar LR, Barber GP, Benet-Pagès A, Casper J, Clawson H, Diekhans M, et al. The UCSC Genome Browser database: 2023 update. Nucleic Acids Res. 2023 Jan 6;51(D1):D1188–95.

39. Gel B, Serra E. karyoploteR: an R/Bioconductor package to plot customizable genomes displaying arbitrary data. Bioinformatics. 2017 Oct 1;33(19):3088–90.

40. Bolger AM, Lohse M, Usadel B. Trimmomatic: a flexible trimmer for Illumina sequence data. Bioinformatics. 2014 Aug 1;30(15):2114–20.

41. Li H. Aligning sequence reads, clone sequences and assembly contigs with BWA-MEM. 2013.

42. Kuilman T, Velds A, Kemper K, Ranzani M, Bombardelli L, Hoogstraat M, et al. CopywriteR: DNA copy number detection from off-target sequence data. Genome Biol. 2015 Feb 27;16:49.

43. Chen B, Gilbert LA, Cimini BA, Schnitzbauer J, Zhang W, Li GW, et al. Dynamic Imaging of Genomic Loci in Living Human Cells by an Optimized CRISPR/Cas System. Cell. 2013 Dec 19;155(7):1479–91.

44. Ran FA, Hsu PD, Wright J, Agarwala V, Scott DA, Zhang F. Genome engineering using the CRISPR-Cas9 system. Nat Protoc. 2013 Nov;8(11):2281–308.

45. Sack LM, Davoli T, Li MZ, Li Y, Xu Q, Naxerova K, et al. Profound Tissue Specificity in Proliferation Control Underlies Cancer Drivers and Aneuploidy Patterns. Cell. 2018 Apr 5;173(2):499–514.e23.

46. Pagès H, Aboyoun P, Gentleman R, DebRoy S. Biostrings: Efficient manipulation of biological strings [Internet]. 2022. Available from: https://bioconductor.org/packages/Biostrings

47. Morgan M, Pagès H, Obenchain V, Hayden N. Rsamtools: Binary alignment (BAM), FASTA, variant call (BCF), and tabix file import [Internet]. 2022. Available from: https://bioconductor.org/packages/Rsamtools

48. Reijns MAM. WT1_ancestral_culture [Internet]. SRA; 2021 [cited 2023 Jan 18]. Available from: https://www.ncbi.nlm.nih.gov/sra/?term=ERR7477340

49. GATK [Internet]. [cited 2023 Mar 21]. Available from: https://gatk.broadinstitute.org/hc/en-us

50. Knaus BJ, Grünwald NJ. vcfr: a package to manipulate and visualize variant call format data in R. Molecular Ecology Resources. 2017;17(1):44–53.

51. Delignette-Muller ML, Dutang C. fitdistrplus: An R Package for Fitting Distributions. Journal of Statistical Software. 2015 Mar 20;64:1–34.

52. McCarthy DJ, Campbell KR, Lun ATL, Wills QF. Scater: pre-processing, quality control, normalization and visualization of single-cell RNA-seq data in R. Bioinformatics. 2017 Apr 15;33(8):1179–86.

53. Mays J, Davoli T. ID 950110 - BioProject - NCBI [Internet]. 2023 [cited 2023 Sep 12]. Available from: https://www.ncbi.nlm.nih.gov/bioproject/950110

54. Mays JC. karyotapR: CNV Analysis for Tapestri [Internet]. 2023 [cited 2023 Sep 8]. Available from: https://github.com/joeymays/karyotapR

55. Mays J. joeymays/karyotapR: Version 0.1 [Internet]. Zenodo; 2023 [cited 2023 Sep 8]. Available from: https://zenodo.org/record/8305561

56. Mays J. KaryoTap Publication Codebook [Internet]. 2023 [cited 2023 Sep 8]. Available from: https://github.com/joeymays/karyotap-publication

57. Mays J, Davoli T. KaryoTap Enables Aneuploidy Detection in Thousands of Single Human Cells [Internet]. Zenodo; 2023 [cited 2023 Sep 8]. Available from: https://zenodo.org/record/8305841

58. Davoli T, Xu AW, Mengwasser KE, Sack LM, Yoon JC, Park PJ, Elledge SJ. Cumulative haploinsufficiency and triplosensitivity drive aneuploidy patterns and shape the cancer genome. Cell. 2013 Nov 21;155(5):948–62.

59. Shih J, Sarmashghi S, Zhakula-Kostadinova N, Zhang S, Georgis Y, Hoyt SJ, Cucinotta CE, Schwartz C, Volkova O, Nurk S, Meredith M, Lajoie BR, Eng K, Shumate A, Gershman A, Rhie A, Soto DC, Perez GE, Miga KH, O’Neill RJ, Eichler EE, Sullivan BA, Phillippy AM, Mao JH, Pevzner PA, Alexandrov LB, Bafna V. Cancer aneuploid genomes exhibit distinctive mutational signatures. Nature. 2023 Jun;618(7965):746–751.

60. Navin N, Kendall J, Troge J, Andrews P, Rodgers L, McIndoo J, Cook K, Stepansky A, Levy D, Esposito D, Muthuswamy L, Krasnitz A, McCombie WR, Hicks J, Wigler M. Tumour evolution inferred by single-cell sequencing. Nature. 2011 Apr;472(7341):90–4.

61. Baslan T, Kendall J, Rodgers L, Cox H, Riggs M, Stepansky A, Troge J, Ravi K, Esposito D, Lakshmi B, Wigler M, Navin N, Hicks J. Genome-wide copy number analysis of single cells. Nat Protoc. 2012 May;7(6):1024–41.

62. Mahlamäki EH, Bärlund M, Tanner M, Gorunova L, Höglund M, Karhu R, Kallioniemi A. Frequent amplification of 8q24, 11q, 17q, and 20q-specific genes in pancreatic cancer. Genes Chromosomes Cancer. 2002 Oct;35(4):353-8.

63. Hodgson JG, Chin K, Collins C, Gray JW. Genome amplification of chromosome 20 in breast cancer. Breast Cancer Res Treat. 2003 Apr;78(3):337–45.

64. Shoshani O, Brunner SF, Yaeger R, Ly P, Nechemia-Arbely Y, Kim DH, Fang R, Castillon GA, Yu M, Li JSZ, Sun Y, Ellisman MH, Ren B, Pevzner PA, Dixon JR, Misteli T, Stephens PJ, Cleveland DW. Chromothripsis drives the evolution of gene amplification in cancer. Nature. 2021 Mar;591(7848):137–141.

65. Dumont M, Gamba R, Gestraud P, Klaasen S, Worrall JT, De Vries SG, Boudreau V, Salinas-Luypaert C, Maddox PS, Lens SM, Kops GJ, McClelland SE, Miga KH, Fachinetti D. Human chromosome-specific aneuploidy is influenced by DNA-dependent centromeric features. EMBO J. 2020 Jan 15;39(2):e102924.

66. Hwang KT, Han W, Cho J, Lee JW, Ko E, Kim EK, Jung SY, Jeong EM, Bae JY, Kang JJ, Yang SJ, Kim SW, Noh DY. Genomic copy number alterations as predictive markers of systemic recurrence in breast cancer. Int J Cancer. 2008 Oct;123(8):1807–15.

67. Sack LM, Davoli T, Li MZ, Li Y, Xu Q, Naxerova K, Wooten EC, Bernardi RJ, Martin TD, Chen T, Leng K, Liang AC, Scorsone KA, Westbrook TF, Wong KK, Elledge SJ. Profound Tissue Specificity in Proliferation Control Underlies Cancer Drivers and Aneuploidy Patterns. Cell. 2018 Apr;173(2):499–514.e23.

68. Rama Ballesteros AR, Hernández R, Perazzoli G, Cabeza L, Melguizo C, Vélez C, Prados J. Specific driving of the suicide E gene by the CEA promoter enhances the effects of paclitaxel in lung cancer. Cancer Gene Ther. 2020 Sep;27(9):657–668.

69. Klaasen SJ, Truong MA, van Jaarsveld RH, Koprivec I, Štimac V, de Vries SG, Risteski P, Kodba S, Vukušić K, de Luca KL, Marques JF, Gerrits EM, Bakker B, Foijer F, Kind J, Tolić IM, Lens SMA, Kops GJPL. Nuclear chromosome locations dictate segregation error frequencies. Nature. 2022 Jul;607(7919):604–609.

70. Lin Y, Wang J, Wang K, Bai S, Thennavan A, Wei R, Yan Y, Li J, Elgamal H, Sei E, Casasent A, Rao M, Tang C, Multani AS, Ma J, Montalvan J, Nagi C, Winocour S, Lim B, Thompson A, Navin N. Normal breast tissues harbour rare populations of aneuploid epithelial cells. Nature. 2024 Dec;636(8043):663–670.

71. Ciani Y, Fedrizzi T, Prandi D, Lorenzin F, Locallo A, Gasperini P, Franceschini GM, Benelli M, Elemento O, Fava LL, Inga A, Demichelis F. Allele-specific genomic data elucidate the role of somatic gain and copy-number neutral loss of heterozygosity in cancer. Cell Syst. 2022 Feb;13(2):183–193.e7.

72. Watson EV, Lee JJ-K, Gulhan DC, Melloni GEM, Venev SV, Magesh RY, Frederick A, Chiba K, Wooten EC, Naxerova K, Dekker J, Park PJ, Elledge SJ. Chromosome evolution screens recapitulate tissue-specific tumor aneuploidy patterns. Nat Genet. 2024 May;56(5):900–912.

73. Jubran J, Slutsky R, Rozenblum N, Rokach L, Ben-David U, Yeger-Lotem E. Machine-learning analysis reveals an important role for negative selection in shaping cancer aneuploidy landscapes. Genome Biol. 2024 Apr 15;25(1):95.

74. Santaguida S, Tighe A, D’Alise AM, Taylor SS, Musacchio A. Dissecting the role of MPS1 in chromosome biorientation and the spindle checkpoint through the small molecule inhibitor reversine. J Cell Biol. 2010 Jul 26;190(1):73–87

75. Williams MJ, Oliphant MUJ, Chen Z, Fowler JC, McFarland JM, Ben-David U. Luminal breast epithelial cells of BRCA1 or BRCA2 mutation carriers and noncarriers harbor common breast cancer copy number alterations. Nat Genet. 2024 Dec;56(12):2753–2762.

76. Davoli T, Uno H, Wooten EC, Elledge SJ. Tumor aneuploidy correlates with markers of immune evasion and with reduced response to immunotherapy. Science. 2017 Jan 20;355(6322):eaaf8399.

77. Alfieri F, Senger G, Oliveto G, Kogenaru M, Davoli T, Schaefer M. Explainable machine learning reveals forces acting on amplifications and deletions across cancer genomes. Back-to-back submission, 2025.

78. Karlsson K, Przybilla MJ, Kotler E, Xu H, Ma Z, Liu K, Curtis C. Deterministic evolution and stringent selection during preneoplasia. Nature. 2023 Jun;618(7937):383–393

